# Psychotic-like experiences, polygenic risk scores for schizophrenia and structural properties of the salience, default mode and central-executive networks in healthy participants from UK Biobank

**DOI:** 10.1101/729921

**Authors:** C. Alloza, M. Blesa-Cábez, M.E. Bastin, J.W. Madole, C.R. Buchanan, J. Janssen, J. Gibson, I.J. Deary, E.M. Tucker-Drob, H.C. Whalley, C. Arango, A.M McIntosh, S.R Cox, S.M Lawrie

**Affiliations:** Department of Psychiatry, The University of Edinburgh, Edinburgh, United Kingdom; Department of Child and Adolescent Psychiatry, Institute of Psychiatry and Mental Health, Hospital General Universitario Gregorio Marañón, Madrid, Spain; Instituto de Investigación Sanitaria Gregorio Marañón (IiSGM), Madrid, Spain; Ciber del Area de Salud Mental (CIBERSAM), Madrid, Spain; MRC Centre for Reproductive Health, The University of Edinburgh, Edinburgh, United Kingdom; Centre for Clinical Brain Sciences, The University of Edinburgh, Edinburgh, United Kingdom; Centre for Cognitive Ageing and Cognitive Epidemiology, Department of Psychology, The University of Edinburgh, Edinburgh, United Kingdom; Department of Psychology, University of Texas at Austin, Austin, USA; Scottish Imaging Network: A Platform for Scientific Excellence (SINAPSE); School of Medicine, Universidad Complutense, Madrid, Spain

**Keywords:** Brain structure, Schizophrenia, Polygenic risk, Psychotic-like experiences, Brain networks, Structural Equation Modelling

## Abstract

Schizophrenia is a highly heritable disorder with considerable phenotypic heterogeneity. Hallmark psychotic symptoms can be considered as existing on a continuum from non-clinical to clinical populations. Assessing genetic risk and psychotic-like experiences (PLEs) in non-clinical populations and their associated neurobiological underpinnings can offer valuable insights into symptom-associated brain mechanisms without the potential confounds of the effects of schizophrenia and its treatment. We leveraged a large population-based cohort (UKBiobank) including information on PLEs, polygenic risk scores for schizophrenia (PRS_SZ_) and multi-modal brain imaging in combination with network neuroscience. Morphometric (cortical thickness, volume) and water diffusion (fractional anisotropy) properties of the regions and pathways belonging to the salience, default-mode and central-executive networks were computed. We hypothesized that these anatomical concomitants of functional dysconnectivity would be negatively associated with PRS_SZ_ and PLEs. PRS_SZ_ was significantly associated with a latent measure of cortical thickness across the salience network (r = −0.069, *p* = 0.010) and PLEs showed a number of significant associations with properties of the salience and default mode networks (involving the insular cortex, supramarginal gyrus and pars orbitalis, *p_FDR_* < 0.050); with the cortical thickness of the insula largely mediating the relationship between PRS_SZ_ and auditory hallucinations. These results are consistent with the hypothesis that higher genetic liability for schizophrenia is related to subtle disruptions in brain structure and predisposes to PLEs even among healthy participants. In addition, our study suggests that networks engaged during auditory hallucinations show structural associations with PLEs in the general population.

## Introduction

Schizophrenia is associated with a range of alterations in brain structure and function [1–4], some of which are related to a family history or specific genetic risk factors. Consistent relationships between delusions, hallucinations and brain structure and function have, however, proved elusive – potentially because of power issues in relatively small clinical samples and confounds such as antipsychotic drug exposure and substance abuse. Psychotic-like experiences (PLEs) of lesser severity are present not only in patients but also in 5-8% of the general population [5], and to some extent predict transition to psychiatric disorders among those with higher PLEs [6]; with cohort studies supporting a continuity between subclinical and clinically significant psychotic symptoms [7, 8]. A recent study found shared genetic aetiology between PLEs and several psychiatric and neurodevelopmental disorders, indicating that PLEs may be related to a general risk for mental health disorders [9]. Studies of the relationship between PLEs and brain imaging metrics have, however, been scarce and characterised by small sample sizes (with N ranging from 25 for auditory hallucinations to 76 for any PLEs [10–13]). Functionally, PLEs have been associated with altered brain dynamics, in particular with default-mode hypoconnectivity [11, 14]. Nonetheless, such evidence validates the study of the pathophysiology of these clinical phenotypes in non-clinical populations [15], with large studies from the general population also offering increased power to detect such effects.

Aberrant functioning and organization of the salience network, default mode network (DMN) and central-executive network (CEN) are core features of several psychiatric and neurological disorders [16]; with patients with schizophrenia showing structural and functional impairments in all three networks [17]. The salience network includes the insula and anterior cingulate cortex and limbic regions, and is involved in the identification of biological and behaviourally relevant stimuli and the subsequent coordination of neural resources to guide flexible behaviour [18, 19]. Aberrant intrinsic functional connectivity of the salience network has been observed in schizophrenia [20, 21] and in individuals at clinical high risk for psychosis [22], and is posited to underlie persecutory delusions in particular [23]. The DMN is a distributed system of fronto-temporal-parietal cortex that is activated during passive cognitive states and deactivates during several cognitive tasks [24]. In schizophrenia, the DMN is overactive with significant correlations between the activity of subregions of the DMN and positive symptoms [25]; although the evidence is somewhat inconsistent [25–27]. The CEN is a frontoparietal system coactivating the dorsolateral prefrontal cortex and the posterior parietal cortex during several cognitive tasks [28]. Increased functional connectivity between the DMN and CEN has been linked to hallucinations in patients with schizophrenia [20].

Schizophrenia is highly polygenic, with many common alleles of small effect, and increasing numbers of genome-wide significant loci have been identified as sample sizes increase [29–31]. Summary statistics from large-scale GWAS allow the degree of genetic liability for a heritable trait to be estimated in healthy subjects outside the population in which the original GWAS was conducted [32, 33]. Only a small number of studies have analysed the relationship between polygenic risk score for schizophrenia (PRS_SZ_) and neuroimaging biomarkers in healthy samples [32–37] but some of these associations map on to regions likely to be involved in the generation of psychotic symptoms and PLEs.

Thus, in this study we investigated how PRS_SZ_ relate to neuroanatomical properties of the salience network, DMN and CEN, and thence to PLEs; in addition to formally testing whether the association between PRS_SZ_ and PLEs was mediated by brain structure. We computed the trajectories of water diffusion MRI parameters (using fractional anisotropy; FA), cortical thickness (CT) and grey matter volume (GMV) of the regions involved in these networks in a large sample of healthy participants from UKBiobank in whom any such associations would not be confounded by illness-associated factors. We employed a novel approach based on ROI-ROI analysis (derived from connectome processing) which extends our previous *a priori* network selection methods [38, 39], allowing a much finer-grained network approach than using other techniques which quantify white matter connectivity without direct subject-specific linkage to cortical or subcortical regions. In addition, previous studies have suggested that schizophrenia may be accompanied by accelerated ageing [40], indicating for instance, significant declines in white matter coherence more than twice that of age-matched controls [41]. Therefore, to examine possible accelerated brain ageing in brain structure, we included an interaction term between age at MRI scanning and PRS_SZ_ in all analyses.

## Methods

### Participants

UKBiobank (http://www.ukbiobank.ac.uk/) comprises around 500,000 community-dwelling participants recruited from across the United Kingdom of Great Britain and Northern Ireland between 2006 and 2014. A subset of the participants who were part of the initial recruitment attended for head MRI scanning at an average of around 4 years after the initial visit (all data presented in this analysis were collected on the same scanner). The current study uses the 5k neuroimaging data release (Supplementary Material (SM) Figure 1). UKBiobank received ethical approval from the Research Ethics Committee (reference 11/NW/0382). Those participants who had been admitted to a hospital with a diagnosis of schizophrenia or bipolar disorder with psychotic symptoms were excluded from our analysis. In order to comprehensibly study the PLEs phenotype, additional analyses comprised: (1) the exclusion of participants who had any psychiatric-related admissions (diagnoses in SM Table 1) and (2) the inclusion of the whole sample (N = 3875). The present analyses were conducted as part of UKBiobank application 16124, linked to 4844 and 10279. All participants provided informed consent (http://biobank.ctsu.ox.ac.uk/crystal/field.cgi?id = 200).

**Figure 1.**
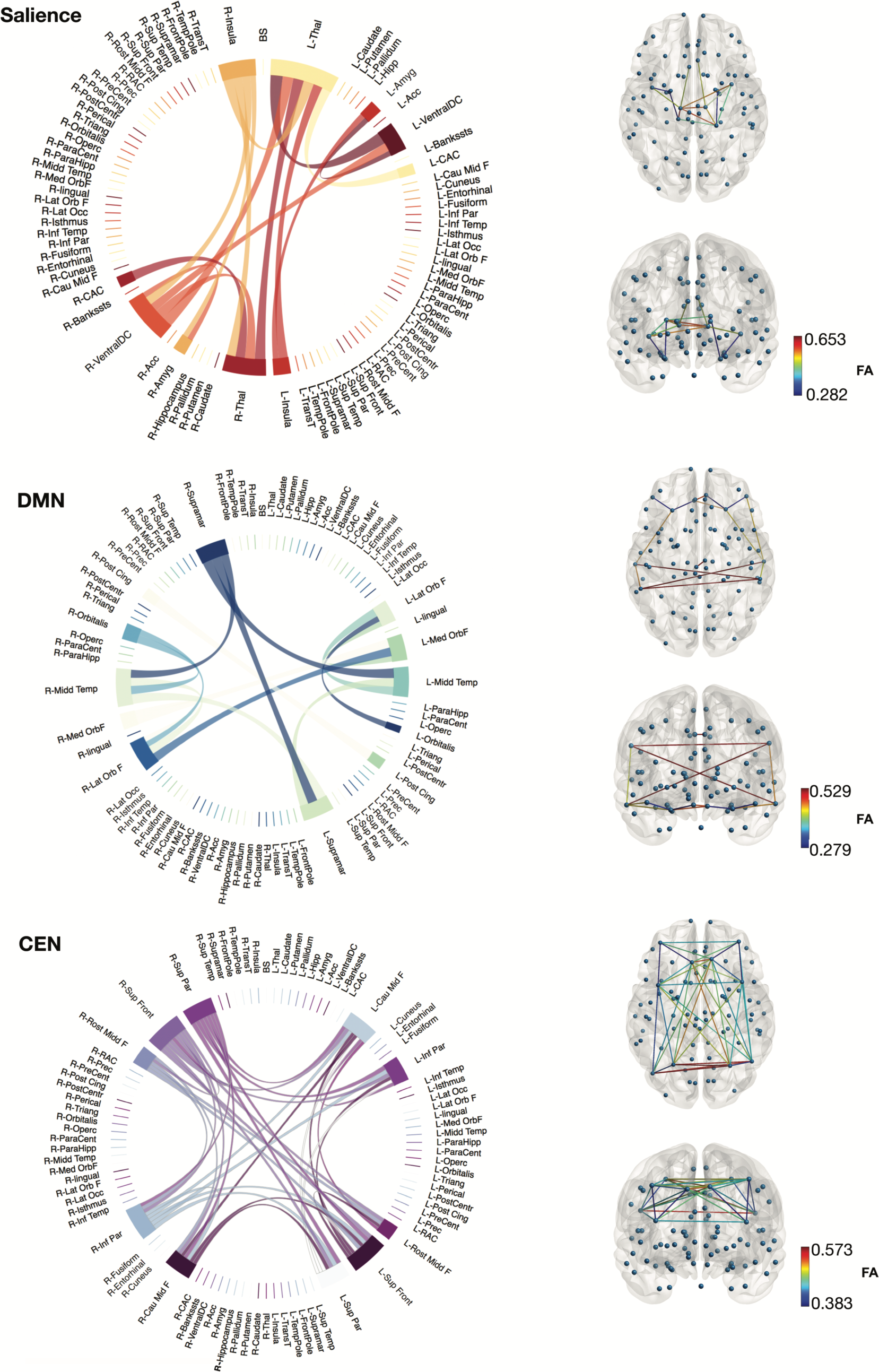
Diagram of nodes of interest and their linking pathways derived from the connectome for the networks of interest. Colours were randomly assigned, and width of the links are proportional to mean FA values across all participants. Axial and coronal views of the networks with colour-coded FA gradient. L: left, R: right. A list of abbreviations is provided in SM Table 2.

**Table 1.** Selected networks and their corresponding nodes.

| Network | Nodes implicated |
| --- | --- |
| Salience | Thalamus, amygdala, VentralDC, caudal anterior cingulate and insula [18, 47] |
| Default mode | Lateral orbitofrontal, medial orbitofrontal, middle temporal, pars orbitalis, posterior cingulate and supramarginal [28, 48] |
| Central-Executive | Caudal middle frontal, inferior parietal, rostral middle frontal, superior frontal and superior parietal [47, 49, 50] |

*Note:* VentralDC = Ventral diencephalon

**Table 2.**
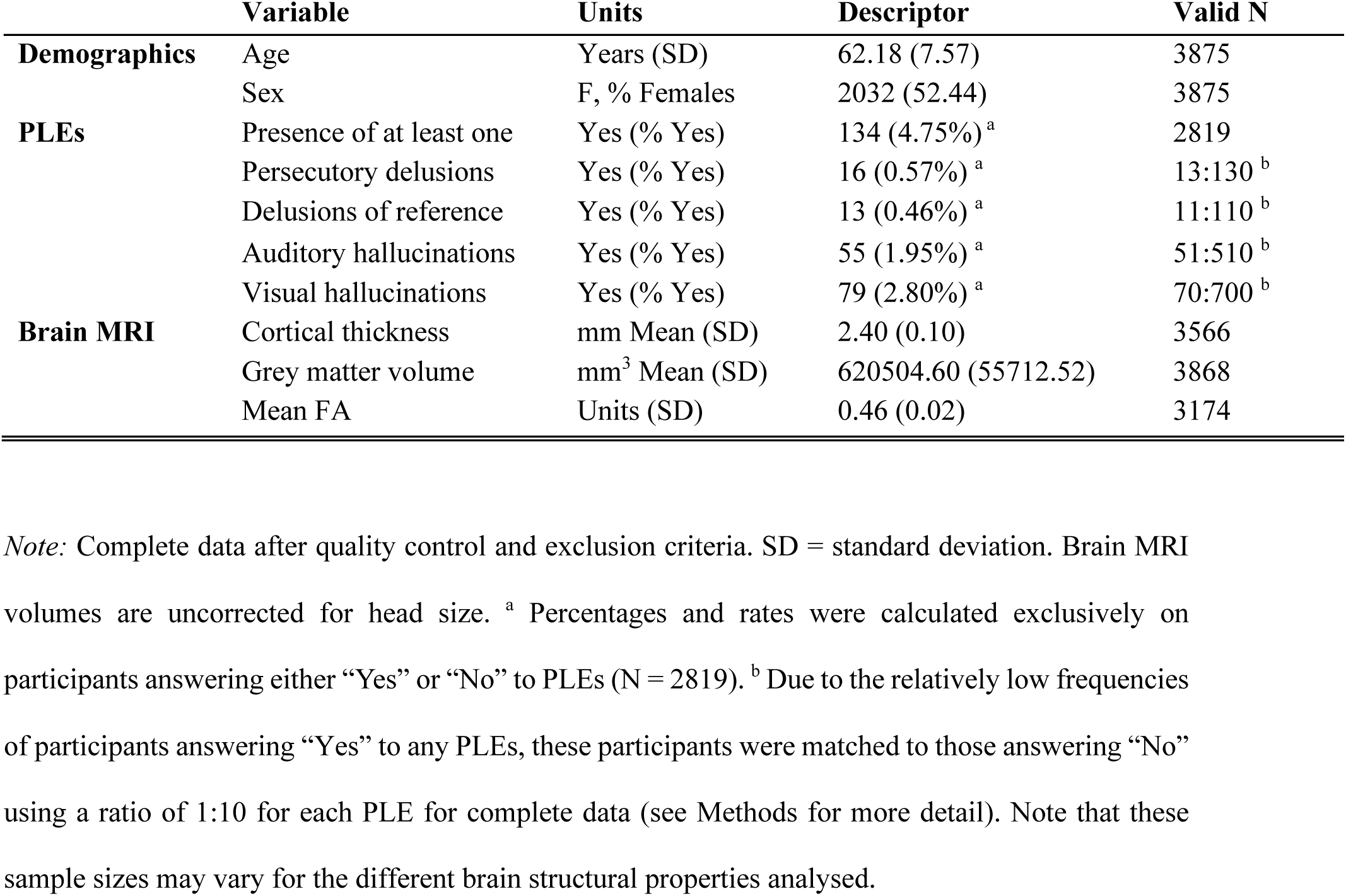
Participants characteristics.

|  | Variable | Units | Descriptor | Valid N |
| --- | --- | --- | --- | --- |
| **Demographics** | Age | Years (SD) | 62.18 (7.57) | 3875 |
|  | Sex | F, % Females | 2032 (52.44) | 3875 |
| **PLEs** | Presence of at least one | Yes (% Yes) | 134 (4.75%)<sup>a</sup> | 2819 |
|  | Persecutory delusions | Yes (% Yes) | 16 (0.57%)<sup>a</sup> | 13:130<sup>b</sup> |
|  | Delusions of reference | Yes (% Yes) | 13 (0.46%)<sup>a</sup> | 11:110<sup>b</sup> |
|  | Auditory hallucinations | Yes (% Yes) | 55 (1.95%)<sup>a</sup> | 51:510<sup>b</sup> |
|  | Visual hallucinations | Yes (% Yes) | 79 (2.80%)<sup>a</sup> | 70:700<sup>b</sup> |
| **Brain MRI** | Cortical thickness | mm Mean (SD) | 2.40 (0.10) | 3566 |
|  | Grey matter volume | mm<sup>3</sup> Mean (SD) | 620504.60 (55712.52) | 3868 |
|  | Mean FA | Units (SD) | 0.46 (0.02) | 3174 |

*Note:* Complete data after quality control and exclusion criteria. SD = standard deviation. Brain MRI volumes are uncorrected for head size. <sup>a</sup> Percentages and rates were calculated exclusively on participants answering either “Yes” or “No” to PLEs (N = 2819). <sup>b</sup> Due to the relatively low frequencies of participants answering “Yes” to any PLEs, these participants were matched to those answering “No” using a ratio of 1:10 for each PLE for complete data (see Methods for more detail). Note that these sample sizes may vary for the different brain structural properties analysed.

### Polygenic risk score calculation

The details of the array design, genotyping, quality control and imputation have been described previously [42]. Quality control included removal of participants based on missingness, relatedness, gender mismatch non-British ancestry and participants based upon overlap in Psychiatric Genomics Consortium (PGC) prediction samples and schizophrenia status. Polygenic profiles were created for schizophrenia in all the genotyped participants using PRSice [43]. PRSice calculates the sum of alleles associated with the phenotype of interest across many genetic loci, weighted by their effect sizes estimated from a genome-wide association study of that phenotype in an independent sample. Before creating the scores, the SNPs with a minor allele frequency <1% were removed and clumping was used to obtain SNPs in linkage equilibrium with an *r*^2^ < 0.25 within a 200 bp window. Five scores were created for each individual using SNPs selected according to the significance of their association with the phenotype at nominal p-value thresholds of 0.01, 0.05, 0.1, 0.5 and 1.0 (all SNPs). Our primary analyses used scores generated from a list of SNPs with a GWAS training set of p ≤ 0.1, as this threshold was shown to explain the most phenotypic variance in the discovery cohort [31] (results for 0.5 and 1.0 thresholds are presented in SM Results 1). Fifteen multidimensional scaling factors (estimated from SNP data) were entered into the models as additional ‘nuisance’ covariates to control for population stratification, along with age and genotyping array.

### Psychotic-like experiences information

Lifetime PLEs were taken from a Mental Health Questionnaire (MHQ; UKBiobank Category: 144) was sent to all participants who provided an email address from July 2016, to July 2017. Responses to the following questions were dichotomised: “Did you ever believe that there was an unjust plot going on to harm you or to have people follow you, and which your family and friends did not believe existed?”, “Did you ever believe that a strange force was trying to communicate directly with you by sending special signs or signals that you could understand but that no one else could understand (for example through the radio or television)?”, “Did you ever hear things that other people said did not exist, like strange voices coming from inside your head talking to you or about you, or voices coming out of the air when there was no one around?”, and “Did you ever see something that wasn’t really there that other people could not see?”. We categorised these PLEs as: persecutory delusions, delusions of reference, and auditory, and visual hallucinations, respectively. This questionnaire explicitly indicated not to include those instances when the participant was dreaming, half-asleep, or under the influences of alcohol or drugs. Moreover, level of distress in relation to PLEs was defined as: “Not distressing at all, it was a positive experience”, “Not distressing, a neutral experience”, “A bit distressing”, “Quite distressing”, and “Very distressing”, and was coded as a continuous variable. Participants that responded “do not know” or “prefer not to answer” were excluded from analyses in all cases.

### Imaging analysis

Full details of the image acquisition and processing can be found on the UKBiobank website (http://biobank.ctsu.ox.ac.uk/crystal/refer.cgi?id=2367), Brain Imaging Documentation (http://biobank.ctsu.ox.ac.uk/crystal/refer.cgi?id=1977) and in [44]. More information regarding scan acquisition and image processing can be found in SM Methods 1 and SM Table 2 for a list of abbreviations of each node. Table 1 and Figure 1 shows the nodes and white matter pathways selected for each of the networks studied here. For the present study, data available for participants who were unrelated, survived the quality control process and had full imaging data available, is provided in Table 2.

### Network construction

For each subject, two networks were constructed: the number of streamline (NOS) network that was created using the number of streamlines connecting each pair of the 85 ROI (network node) pairs from the default FreeSurfer cortical [45] and subcortical regions (Desikan-Killiany atlas, see Supplementary Material Table 2 for a list of abbreviations of each node); and the FA-weighted network that was constructed by recording the mean FA value along streamlines. The endpoint of a streamline was considered to be the first grey matter ROI encountered when tracking from the seed location. In order to reduce the number of spurious connections derived from probabilistic tractography, we applied a consistency-based threshold to the NOS matrices using the numbers of streamlines connecting all 85 ROI and preserving exclusively the top 30% white matter tracts that were most consistent across subjects [46]. This mask was then applied to the FA-weighted connectivity matrices. For each FA-weighted connectivity matrix for the thresholded network, the salience, DMN and CEN masks were applied based on our bilateral nodes of interest. Mean FA was computed using only the non-zero matrix elements.

### Statistical analyses

We undertook an *a priori* network-of-interest (NOI) approach, based on the literature cited above.

#### 1. Linear regressions for individual network components

Initially, we conducted linear regressions between PRS_SZ_ and each node (CT, GMV) and edge (FA) within each of the selected NOIs. Within each model, each morphometric measure was set as the dependent variable, PRS_SZ_ as the independent variable, controlling for age, sex, and the interaction between age and PRS_SZ_.

#### 2. Network analyses

Next, we aimed to test whether associations with PRS_SZ_ were best represented in the data at the network-general level (common pathway), or whether there were additional unique associations with specific network components (common + independent pathways analysis), or whether the associations were simply best described by independent pathways [51]. We did so in a structural equation modelling (SEM) framework. First, we fitted measurement models (i.e. models that relate the latent factor to its manifest variables) to ascertain the degree to which we could describe overall network integrity at the network level; this initial measurement model was a confirmatory factor analysis in which we tried to estimate a single latent construct of global NOI by incorporating altogether grey and white matter metrics pertaining to all nodes within a NOI. However, these models all exhibited poor fit to the data, and the latent measures of the grey and white matter did not correlate significantly.

##### a) Common model analysis

Thus, we opted to construct three measurement models for each NOI in which a latent factor was indicated by all network components separately: CT (derived from the bilateral cortical nodes), GMV (including all bilateral cortical and subcortical nodes), or white matter FA (derived from all pathways connecting nodes within each NOI). These measurement models all fitted the data well except for FA in the DMN and CEN which did not achieve acceptable model fit statistics and thus, were not included in these analyses.

For each model, we then tested associations between PRS_SZ_, age, the interaction PRS_SZ_ *age, and MRI parameters. We did so by fitting a multiple indicators, multiple causes (MIMIC) model [52]; Figure 2 A shows a simplified diagram of the SEM framework. Within the model, each brain imaging measure was adjusted at the manifest level for sex and either whole brain average CT, whole brain total GMV or whole brain mean FA, while PRS_SZ_ was adjusted for sex, population stratification components, and genotyping array. Model fits were assessed using the following indices and cut-offs: CFI, RMSEA and SRMR. We allowed for certain residual correlations between manifest variables in those cases in which their addition improved the model fit significantly. SEM analyses were conducted using the package ‘lavaan’ [53] in R with full-information maximum likelihood estimation to use all data available.

**Figure 2.**
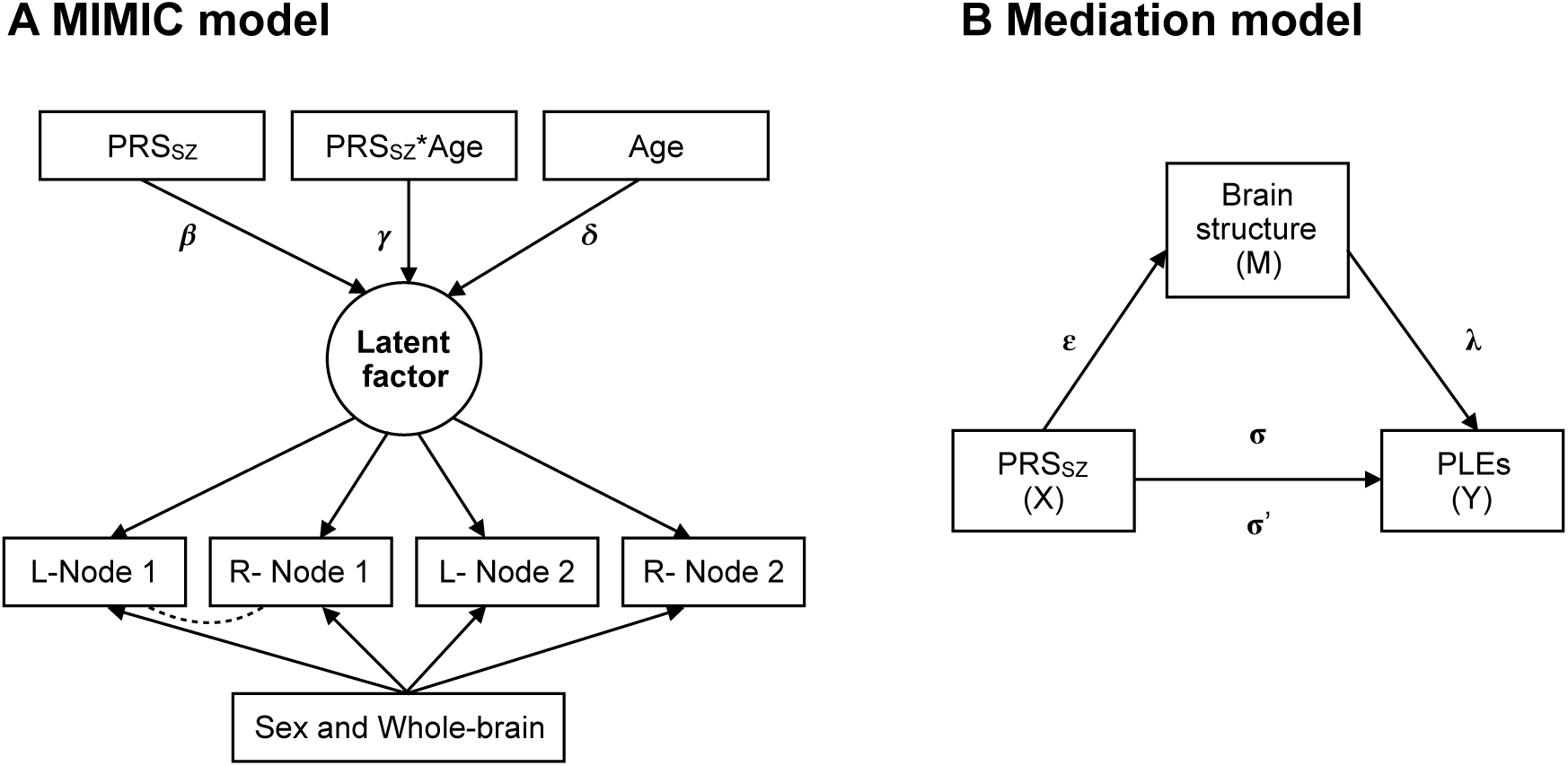
Diagrams of structural equation models (SEM). **A.** Multiple indicators, multiple causes (MIMIC) model [52] for neurostructural properties of each network. A separate model was applied to FA, grey matter thickness (CT) and grey matter volume (GMV). From each individual bilateral node (L: left; R: right) or pathway, a latent score was calculated for FA, CT, GMV controlling for sex and whole-brain structural properties at the manifest level (i.e: whole brain FA/CT/GMV). Relation between FA/CT/GMV and PRS_SZ_ is indicated by path β; path γ represents the association between the interaction of age and PRS_SZ_ and FA/CT/GM factors; path δ represents the association between age and the latent factor. PRS_SZ_ was corrected for sex and population stratification (paths not shown). The dashed line represents a possible residual correlation between nodes. **B.** Path diagram of mediation model, where the ε coefficient represents the regression of X on M, λ coefficient the regression of M on Y and σ coefficient the direct path of X on Y. The product of the ε and λ coefficients describes the indirect path of X on Y through M (σ’).

##### b) MIMIC common + independent pathways analysis

To assess whether associations between PRS_SZ_ and latent constructs of network integrity were best characterized at the network-general level, or whether there were additional associations with particular brain regions beyond this, we ran a common plus independent pathways analysis [51] (see details in SM Methods 2).

#### 3. PLEs

Due to the low percentage of participants who answer “Yes” to experiencing PLEs (see Table 2), we first matched those participants to those who answered “No” by age and sex with a ratio 1:10 using a propensity score method in the ‘MatchIt’ package [54]. Linear regressions were then calculated using brain regions and each PLEs as the response variable, covarying for either whole brain total volume (for node’s volume; to control for differences in whole-brain volume across participants) or average CT (for node’s CT; to control for differences in whole-brain CT across participants), age at MRI and sex. Raincloud plots [55] were used to plot group differences between participants with PLEs and their matched controls.

#### 4. Mediation analyses

We further sought to formally test whether the association between PRS_SZ_ and PLEs was mediated by brain structure (at the manifest level). We did this in the form of a mediation model using the ‘lavaan’ package [53] with full-information maximum likelihood estimation, confidence intervals were reported (CI; constructed using 1000 bootstraps). This analysis was performed only in those cases where the path between brain structure (M) and PLEs (Y) was significant (*p*FDR < 0.050); mediation was observed when the change from σ to σ’ was statistically significant (confidence intervals did not include zero). Within the model, each brain region was adjusted for age, sex and either average CT (CT analysis) or whole brain total volume (GMV analysis). Figure 2 B shows a simplified diagram of the mediation framework.

#### 5. Additional analyses

In order to determine whether the signal could be driven by any complication associated with any psychiatric disorder beyond a diagnosis of psychosis (i.e.: distress, dysfunction, co-morbidities, medication, unhealthy lifestyles, etc.), we also performed an analysis that excluded all participants with any psychiatric diagnoses from the PLEs analysis (SM Results 2). Due to the possible clinical significance of level of distress in relation to PLEs, we first computed logistic regressions with PLEs as the response variable and level of distress, age and sex as predictor variables. For those significant associations, we tested whether the level of distress caused by PLEs could be mediating the relationship between brain structure and PLEs (SM Results 3). We also tested the hypothesis that increased number of PLEs (i.e individual sum of these four types of PLEs) was associated with a higher level of distress [56] (SM Results 4).

Analyses were performed in R (https://www.r-project.org) and standardised betas were reported throughout. For each section/model of statistical analysis, significance (*p*) values (α = 0.050) were corrected for multiple comparisons using false discovery rate (FDR) [57].

## Results

Participant characteristics are presented in Table 2, and PLEs prevalence in SM Table 3. Mean values for CT and GMV for each node (according to the Desikan-Killiany atlas [45]) are shown in SM Figure 2. All SEM models showed acceptable fit to the data (fit indices are shown in SM Table 4).

### PRS_SZ_ ANALYSES

#### Salience Network

##### Linear regressions for individual network components

There were no significant associations between cortical and subcortical GMV and PRS_SZ_ (*p_FDR_* > 0.050). For CT, nominally significant negative associations were found between the right and left insula and PRS_SZ_ (β = −0.046, SE = 0.019, *p_FDR_* = 0.050, β = −0.039, SE = 0.018, *p_FDR_* = 0.065, respectively). Interactions between age and PRS_SZ_ were significant for CT of the right caudal anterior cingulate (CAC) (β = −0.054, SE = 0.021, *p_FDR_* = 0.040). There were no significant associations between FA and PRS_SZ_ (*p_FDR_* > 0.050).

##### MIMIC common + independent pathways analysis

The association between the latent factor for GMV and PRS_SZ_ was not significant (r = −0.026, SE = 0.014, *p* = 0.242). Allowing for a direct effect of PRS_SZ_ on left thalamus GMV significantly improved model fit (χ^2^[1] > 4.917, p < 0.028; independent pathway estimates: β = 0.017, *p* = 0.032). A direct effect of right CAC GMV on PRS_SZ_*age (β = −0.035, *p* = 0.088) – the second independent pathway added – significantly improved model fit (χ^2^[1] = 5.893, *p* = 0.015). Results are shown in Figure 3A. In addition to the association between PRS_SZ_ and the latent factor of CT (r = −0.069, SE = 0.015, *p* = 0.010), an independent pathway emerged from PRS_SZ_*age to right CAC CT (β = −0.043, *p* = 0.031), which improved the model fit (χ^2^[1] = 4.606, *p* = 0.03; Figure 3B). There was no significant association between the latent factor for FA and either PRS_SZ_ or PRS_SZ_*age interaction (*p* > 0.050).

**Figure 3.**
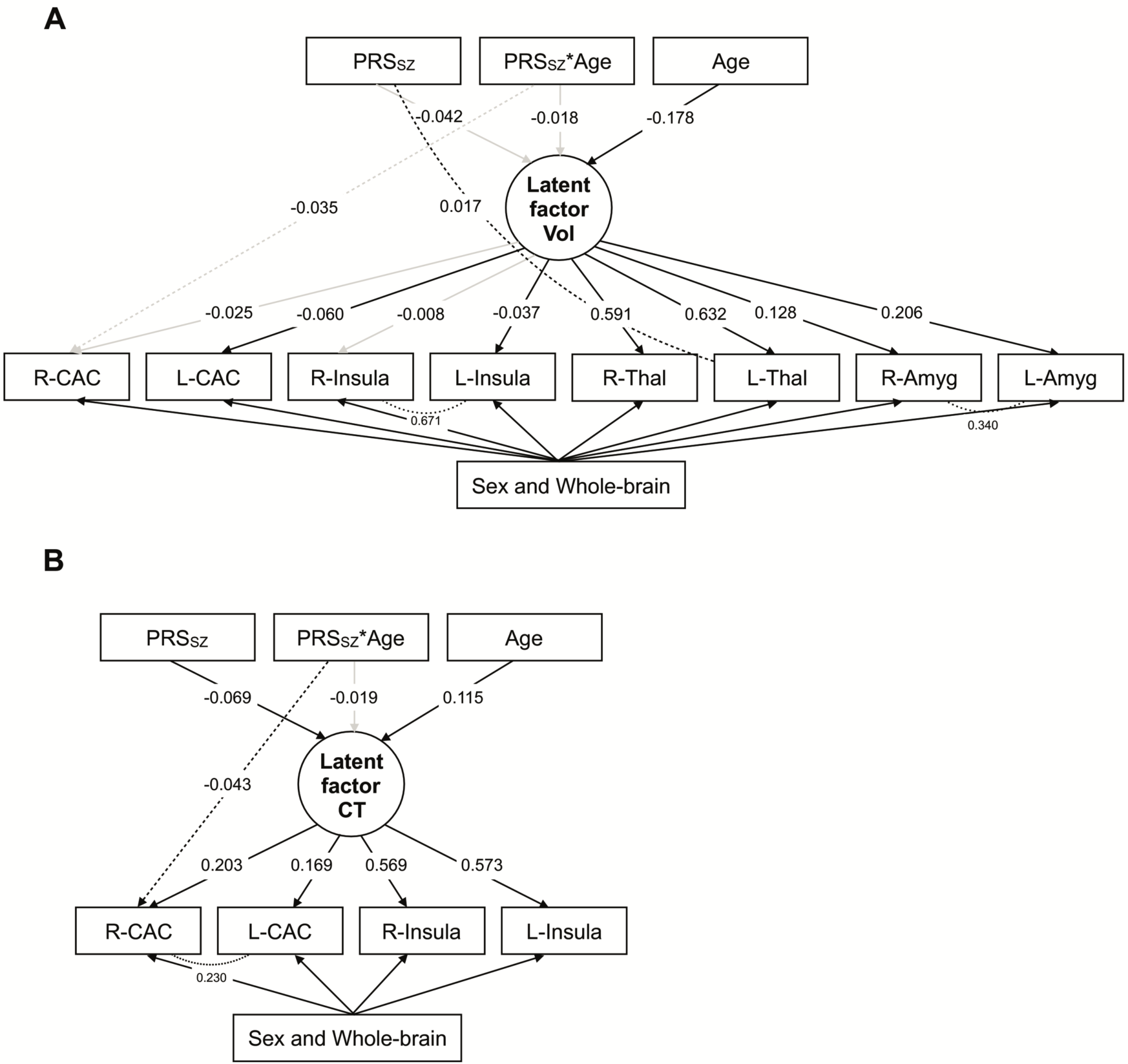
Diagrams of MIMIC and independent pathway models of the salience network. **A.** Common + independent pathways model for latent factor of grey matter volume. **B.** Common + independent pathways model for latent factor of grey matter cortical thickness. Dashed lines represent independent pathways while dotted lines residual correlations between nodes. Black lines represent statistically significant pathways (*p* < 0.050) while grey lines non-significant pathways (*p* > 0.050).

#### Default Mode Network

##### Linear regressions for individual network components

There were no significant associations between GMV, CT or FA and PRS_SZ_ or the interaction between PRS_SZ_*age (p_FDR_ > 0.050).

##### MIMIC common + independent pathways analysis

The associations between the latent factors for GMV and CT and PRS_SZ_ were not significant (r_GMV_ = - 0.040, SE = 0.012, p = 0.111; r_CT_ = −0.028, SE = 0.015, p = 0.273). An independent pathway emerged from PRS_SZ_*age to right supramarginal CT (χ^2^[1] > 5.705, p < 0.018; independent pathway estimate: β = −0.035, p = 0.008). Interactions between PRS_SZ_*age were not significantly associated with the latent constructs (p > 0.050).

#### Central Executive Network

##### Linear regressions for individual network components

No significant associations were found between the GMV, CT or FA and PRS_SZ_, or the interaction between PRS_SZ_*age (p > 0.050).

##### MIMIC common + independent pathways analysis

The associations between the latent factors for grey matter and PRS_SZ_ were not significant (p > 0.050). For GMV, allowing for a direct effect of PRS_SZ_ on right inferior parietal GMV (β = 0.039, p = 0.004) resulted in significant improvement in model fit (χ^2^[1] > 7.741, p < 0.006). For CT, an independent pathway emerged from PRS_SZ_*age to right superior frontal CT (χ^2^[1] > 7.512, p < 0.007; independent pathway estimate: β = 0.028, p =0.002) and right inferior parietal CT (χ^2^[1] > 4.314, p < 0.039; independent pathway estimate: β = −0.024, p =0.034). Interactions between PRS_SZ_*age were not significantly associated with any latent construct (p_FDR_ > 0.050).

### PSYCHOTIC-LIKE EXPERIENCES

Table 2 shows the frequencies and percentages of respondents with PLEs, SM Table 3 shows sample characteristics for participants reporting PLEs versus non-PLEs. Figure 4A shows raincloud plots representing differences in CT and GMV between participants reporting PLEs and their matched controls.

**Figure 4:**
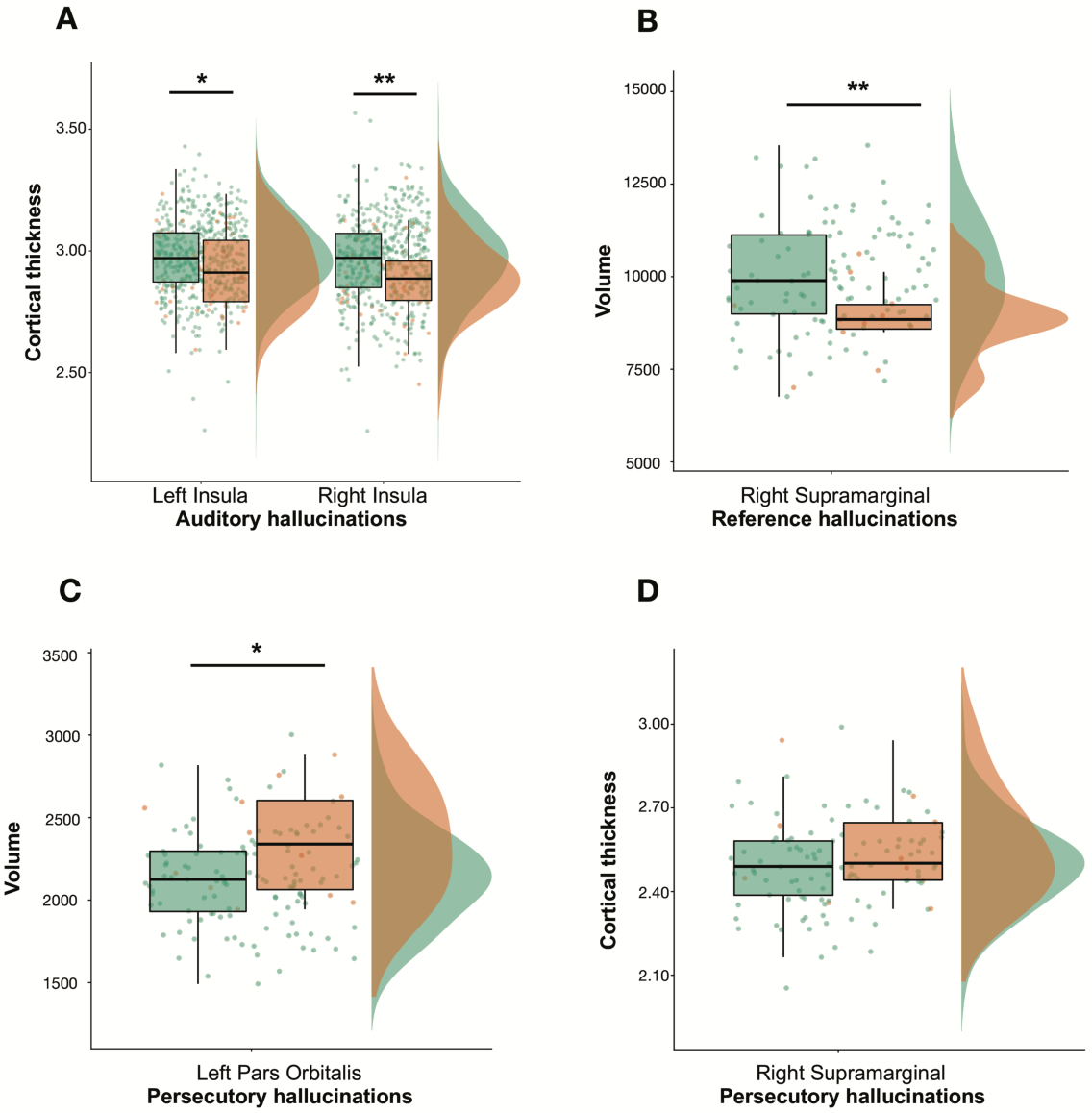
Raincloud plots representing differences in cortical thickness (CT) and volume (GMV) between participants reporting PLEs (orange) and matched controls (green). **A** Bilateral CT of the insula and auditory hallucinations. **B** GMV of right supramarginal gyrus and reference hallucinations. **C** GMV of the left pars orbitalis and persecutory hallucinations. **D** CT of the right supramarginal gyrus and persecutory hallucinations. *Note*: CT is measured in cm and GMV in cm^3^. Asterisks represent significant differences between groups (*: *p <* 0.05, **: *p* < 0.01).

**Figure 5:**
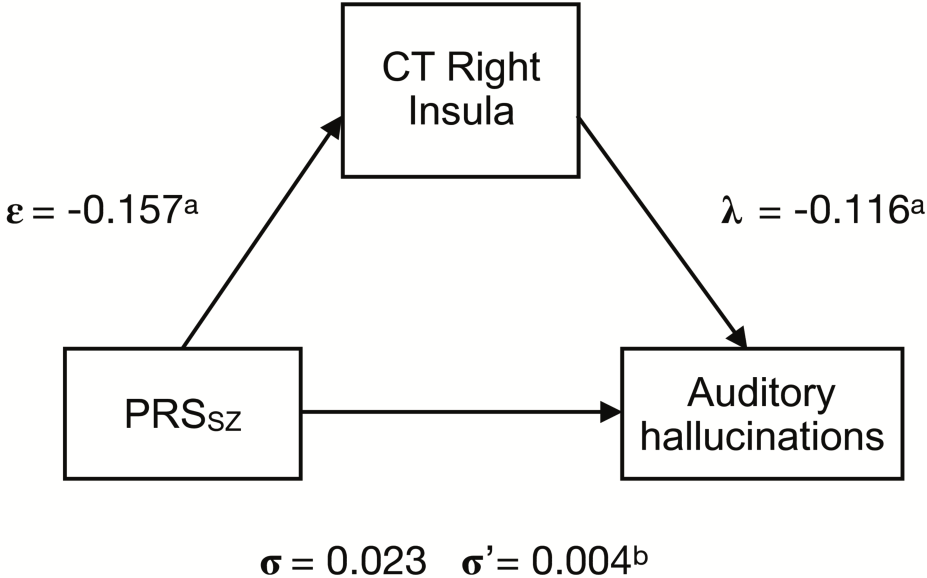
Path diagram of the mediation model, where the ε coefficient represents the coefficient of regressions of PRS_SZ_ on cortical thickness (CT) of the right insula, and λ the coefficient of the regression of CT of the right insula on auditory hallucinations and σ coefficient of the direct path of PRS_SZ_ and auditory hallucinations. Coefficient of σ’ represents the indirect path of PRS_SZ_ on auditory hallucinations through CT of the right insula. *Note:* ^a^ *p* values < 0.01, ^b^ Confidence intervals not including zero.

Within the salience network, significant negative associations were found between the CT of the right and left insula and auditory hallucinations (N_cases_ = 51, N_controls_ = 510; β = −0.114, SE = 0.035, p_FDR_ = 0.004 and β = −0.083, SE = 0.036, p_FDR_ = 0.045, respectively). Associations between total number of PLEs and CT/GMV of salience network nodes were all non-significant.

For the default mode network, a significant negative association was found between the GMV of the right supramarginal gyrus and delusions of reference (N_cases_ = 11, N_controls_ = 110; β = −0.195, SE = 0.061, p_FDR_ = 0.022) and between persecutory delusions and GMV of the left pars orbitalis (N_cases_ = 12, N_controls_ = 120; β = 0.219, SE = 0.069, p_FDR_ = 0.021). For persecutory delusions, there was a positive association with the CT of the right supramarginal gyrus (N_cases_ = 13, N_controls_ = 130; β = 0.177, SE = 0.048, p_FDR_ = 0.003). We did not find any significant association between total number of PLEs and GMV/CT of nodes. No significant associations were found between CEN’s nodes and PLEs (p_FDR_ > 0.050). The overall PLE findings are summarised in Supplementary Table 5.

#### Mediation analyses

The mediation models using PRS_SZ_ and mediators met the criteria for a close-fitting model (RMSEA = 0, CFI = 1, SRMR < 0.05). We examined the hypothesis that higher PRS_SZ_ was associated with symptom severity via the brain structures significantly associated with PLEs in the salience and DMN. First, we tested the association between PRS_SZ_ and total number of PLEs in the whole UKBiobank sample, independently of diagnosis (N_PRS_ = 302,575, N_PLEs_ = 157,305, with N = 7803 answering “Yes” to experiencing at least one PLEs). However, we did not find any significant association (β = 0.020, SE = 0.018, *p* = 0.283). We found that the linear association between szPGRS and auditory hallucinations was significantly mediated by CT of the right insula (from σ = 0.023 to σ’ = 0.004, CI [0.020, 0.127] with the right insular cortex mediating 82.6% of the association between szPGRS and auditory hallucinations). These results were consistent across all szPGRS thresholds (SM Results 5). No significant mediations were observed for any other ROIs.

## Discussion

The present study aimed to investigate different structural properties of the salience, DMN and CEN networks in relation to PRS_SZ_ and PLEs in a large population-based cohort of healthy participants. This is, to our knowledge, the first study to investigate these polygenic-MRI associations using the structural properties of brain networks from a connectome perspective, where the properties of estimated connections that directly link distal cortical and subcortical regions in each individual are examined. At the node level, we found nominally significant associations between thinner right and left insular cortices and higher PRS_SZ_. At the network level we found that higher PRS_SZ_ was associated with reduced CT across the salience network, with some specific associations between PRS_SZ_ and individual brain properties emerging. The findings reported here indicate both global network and domain-specific pathways implicated in schizophrenia, indicating that those individual regions showing an independent significant association with PRS_SZ_ may be more susceptible/resilient to the effect of PRS_SZ_ beyond a latent factor of network integrity. We found several significant associations between PLEs – in particular regarding auditory hallucinations – and structural properties of the salience and DMN; with the right insular cortex largely mediating the association between auditory hallucinations and PRS_SZ_. Impairments in white matter micro-and macrostructure are a common feature in schizophrenia with healthy relatives who are genetically at higher risk of developing schizophrenia also exhibiting impairments in FA [58]. Our results are in accordance to a previous study, reporting non-significant associations between a general factor of FA (including 27 major white matter tracts) and PRS_SZ_ in a previous release of this data (n = 816) [59], though the current approach offers greater brain regional fidelity and statistical power. Despite the apparent alterations in white matter structure in schizophrenia, only a small number of studies have reported significant associations between PRS_SZ_ and white matter in healthy and clinical samples [34, 37, 60]. Significant reductions in global white matter volume in dizygotic twins discordant for schizophrenia have been reported while reductions in global grey matter were exclusively observed in schizophrenia [61]. Thus, the null findings of white matter observed here may partly be due to the aggregation of genetic and environmental risk factors found in affected individuals and their relatives compared with healthy individuals.

### Psychotic-like experiences

The reported prevalence of PLEs in this imaging sample (4.75%) is similar to that in the whole UK Biobank sample (5%)[62], and with previous population studies [63, 64]. We did not find any significant association between PRS_SZ_ and total number of PLEs in the entire UKBiobank sample (N = 308,693), consistent with previous studies investigating this relationship in adolescence [65, 66]. A recent UKBiobank study found a weak association between the presence of any PLEs and PRS_SZ_ at a threshold of P ≤ 0.05 (OR, 1.09; 95% CI, 1.06-1.12; adjusted R2 = 0.001; P = 2.96×10^−11^, N = 500,000); however, as the authors noted, this association may be biased by the possibility that their analysis included sample overlap between UKBiobank and the training sets [9].

The associations between PLEs and insula volume is consistent with its involvement in interoception; this awareness of the body’s internal state comprises emotional responses, complex cognitive states, and the sense of self [67, 68]. Though our sample sizes were modest they are relatively large compared to the previous literature on PLEs and brain imaging in adults [10–13, 69] (for instance, studies on auditory hallucinations with samples of less than 30 participants compared to our sample of 51 experiencing auditory hallucinations and 134 for any PLEs). Moreover, our results are consistent with correlations between psychotic symptoms, activation, volume, and surface area of the insula in clinical [70, 71] and high-risk samples [22]; with hallucinations being correlated with structural aspects of the insula [72]. Our mediation analyses extend these findings by showing the CT of the right insula largely mediating the association between PRS_SZ_ and auditory hallucinations; in accordance with evidence suggesting that predominant right lateralization may discriminate auditory-verbal hallucinations from normal inner speech [73].

The positive correlation between the GMV of the right pars orbitalis – ventral subregion of the inferior frontal gyrus (IFG) – and persecutory delusions is, however, at odds with previous studies reporting reduced GMV of the pars orbitalis in high risk individuals [6, 74] and negative associations between its volume and positive symptoms in clinical samples [75]. We also found that brain structure abnormalities in the supramarginal gyrus were associated with delusions. This is in line with the role of the supramarginal gyrus in processing auditory inputs, especially language [76], and the relation between lower volume and auditory hallucinations in schizophrenia [77]. Moreover, in non-psychotic individuals, auditory hallucinations have been linked to abnormal brain connectivity within the DMN and with auditory cortices (sample of participants experiencing auditory hallucinations of N = 25 and N = 29, respectively [10, 12]). The involvement of right supramarginal, insula and IFG – all of them language-related areas – in PLEs has been extensively documented, indicating that those experiencing auditory hallucinations tend to show impaired speech perception [78] with an engagement of both speech production and reception circuitry [79]. Voices tend to be the dominant content of auditory hallucinations because the human auditory system is tuned to the natural priors of speech [80] and may contribute to the formation of delusions by attributing meaning/agency to the experience, supported by our findings involving the supramarginal gyrus.

### Limitations

We limited our analysis to those networks implicated in previous functional and structural studies, translating those nodes onto a common parcellation scheme. Given issues of validity and comparability across brain atlases, and the implications of this for underlying connectivity [81–83], further research should aim to replicate this study by computing the networks at different levels of node granularity and with different structural properties. We applied a consistency-based thresholding method in an attempt to remove spurious white matter connections, exclusively preserving the top 30% that were most consistent across subjects. Though current evidence indicates that the human brain is likely to exhibit a fairly high degree of sparsity, principles of white matter connectivity in humans are not sufficiently comprehensive (when compared to the mouse [84]) to allow the confident implementation of detailed anatomical priors [85]; some connections may have therefore been pruned erroneously. We investigated whether neurostructural properties of brain networks are associated with PLEs. However, due to the still modest frequencies of PLEs in this sample, results should be interpreted cautiously. A previous study has indicated a potential “healthy volunteer” selection bias in this sample [86], in particular, people with lower socio-economic status, chronic illness, and smokers were under-represented [62]. This suggests that this cohort may not be representative of the sampling population.

## Conclusions

In a large sample of largely healthy participants from UKBiobank a higher genetic liability for schizophrenia was associated with subtle neurostructural differences; in particular, a thinner cortex across the salience network. Beyond this global relationship, some independent paths emerged in each structural network. We also found significant associations between PLEs and the insula, supramarginal gyrus, and pars orbitalis; with the insula largely mediating the association between PRS_SZ_ and auditory hallucinations. Our results in a large healthy sample support previous studies on aberrant activation of language-related areas and the externalization of inner speech phenomena in clinical samples. The results of this research indicate that the study of the non-clinical phenotype represents a valid approach to investigate the pathophysiology of the clinical phenotype and suggest a shared genetic aetiology between the clinical and non-clinical phenotype.

## Acknowledgements

We thank the UK Biobank participants for their participation and the UK Biobank team for their work in collecting, processing and disseminating these data for analysis. This research was conducted, using the UK Biobank Resource under approved project 16124 linked to 4844, in the Department of Psychiatry and Centre for Cognitive Ageing and Cognitive Epidemiology (CCACE) (http://www.ccace.ed.ac.uk) at The University of Edinburgh. SRC, SJR, JWM, MEB, CRB, IJD and EMT-D were supported by a National Institutes of Health (NIH) research grant R01AG054628. EMT-D was also supported by National Institutes of Health (NIH) research grant R01HD083613, and is a member of the Population Research Center at the University of Texas at Austin, which is supported by NIH center grant P2CHD042849. Also supported by the Spanish Ministry of Science, Innovation and Universities. Instituto de Salud Carlos III, co-financed by ERDF Funds from the European Commission, “A way of making Europe”, CIBERSAM. Madrid Regional Government (B2017/BMD-3740 AGES-CM-2), European Union Structural Funds and European Union Seventh Framework Program and H2020 Program; Fundación Familia Alonso, Fundación Alicia Koplowitz and Fundación Mutua Madrileña.

## References

1. Kelly S, Jahanshad N, Zalesky A, Kochunov P, Agartz I, Alloza C, et al. Widespread white matter microstructural differences in schizophrenia across 4322 individuals: results from the ENIGMA Schizophrenia DTI Working Group. Mol Psychiatry. 2017. 17 October 2017. https://doi.org/10.1038/mp.2017.170.

2. Lawrie SM, Abukmeil SS. Brain abnormality in schizophrenia. A systematic and quantitative review of volumetric magnetic resonance imaging studies. Br J Psychiatry J Ment Sci. 1998;172:110–120.

3. van Erp TGM, Walton E, Hibar DP, Schmaal L, Jiang W, Glahn DC, et al. Cortical Brain Abnormalities in 4474 Individuals With Schizophrenia and 5098 Control Subjects via the Enhancing Neuro Imaging Genetics Through Meta Analysis (ENIGMA) Consortium. Biol Psychiatry. 2018. 14 May 2018. https://doi.org/10.1016/j.biopsych.2018.04.023.

4. Wright IC, Rabe-Hesketh S, Woodruff PW, David AS, Murray RM, Bullmore ET. Meta-analysis of regional brain volumes in schizophrenia. Am J Psychiatry. 2000;157:16–25.

5. van Os J, Linscott RJ, Myin-Germeys I, Delespaul P, Krabbendam L. A systematic review and meta-analysis of the psychosis continuum: evidence for a psychosis proneness-persistence-impairment model of psychotic disorder. Psychol Med. 2009;39:179–195.

6. DeRosse P, Nitzburg GC, Ikuta T, Peters BD, Malhotra AK, Szeszko PR. Evidence from structural and diffusion tensor imaging for frontotemporal deficits in psychometric schizotypy. Schizophr Bull. 2015;41:104–114.

7. Cannon M, Caspi A, Moffitt TE, Harrington H, Taylor A, Murray RM, et al. Evidence for early-childhood, pan-developmental impairment specific to schizophreniform disorder: results from a longitudinal birth cohort. Arch Gen Psychiatry. 2002;59:449–456.

8. Hanssen M, Bak M, Bijl R, Vollebergh W, van Os J. The incidence and outcome of subclinical psychotic experiences in the general population. Br J Clin Psychol. 2005;44:181–191.

9. Legge SE, Jones HJ, Kendall KM, Pardiñas AF, Menzies G, Bracher-Smith M, et al. Association of Genetic Liability to Psychotic Experiences With Neuropsychotic Disorders and Traits. JAMA Psychiatry. 2019. 25 September 2019. https://doi.org/10.1001/jamapsychiatry.2019.2508.

10. Diederen KMJ, Neggers SFW, de Weijer AD, van Lutterveld R, Daalman K, Eickhoff SB, et al. Aberrant resting-state connectivity in non-psychotic individuals with auditory hallucinations. Psychol Med. 2013;43:1685–1696.

11. Barber AD, Lindquist MA, DeRosse P, Karlsgodt KH. Dynamic Functional Connectivity States Reflecting Psychotic-like Experiences. Biol Psychiatry Cogn Neurosci Neuroimaging. 2018;3:443–453.

12. van Lutterveld R, Diederen KMJ, Otte WM, Sommer IE. Network analysis of auditory hallucinations in nonpsychotic individuals. Hum Brain Mapp. 2014;35:1436–1445.

13. Orr JM, Turner JA, Mittal VA. Widespread brain dysconnectivity associated with psychotic-like experiences in the general population. NeuroImage Clin. 2014;4:343–351.

14. Satterthwaite TD, Vandekar SN, Wolf DH, Bassett DS, Ruparel K, Shehzad Z, et al. Connectome-Wide Network Analysis of Youth with Psychosis Spectrum Symptoms. Mol Psychiatry. 2015;20:1508–1515.

15. Kelleher I, Cannon M. Psychotic-like experiences in the general population: characterizing a high-risk group for psychosis. Psychol Med. 2011;41:1–6.

16. Menon V. Large-scale brain networks and psychopathology: a unifying triple network model. Trends Cogn Sci. 2011;15:483–506.

17. Palaniyappan L, Mallikarjun P, Joseph V, White TP, Liddle PF. Regional contraction of brain surface area involves three large-scale networks in schizophrenia. Schizophr Res. 2011;129:163–168.

18. Menon V. Salience Network. Brain Mapp., Elsevier; 2015. p. 597–611.

19. Uddin LQ. Salience processing and insular cortical function and dysfunction. Nat Rev Neurosci. 2015;16:55–61.

20. Manoliu A, Riedl V, Zherdin A, Mühlau M, Schwerthöffer D, Scherr M, et al. Aberrant dependence of default mode/central executive network interactions on anterior insular salience network activity in schizophrenia. Schizophr Bull. 2014;40:428–437.

21. Orliac F, Naveau M, Joliot M, Delcroix N, Razafimandimby A, Brazo P, et al. Links among resting-state default-mode network, salience network, and symptomatology in schizophrenia. Schizophr Res. 2013;148:74–80.

22. Wotruba D, Michels L, Buechler R, Metzler S, Theodoridou A, Gerstenberg M, et al. Aberrant coupling within and across the default mode, task-positive, and salience network in subjects at risk for psychosis. Schizophr Bull. 2014;40:1095–1104.

23. Kapur S. Psychosis as a state of aberrant salience: a framework linking biology, phenomenology, and pharmacology in schizophrenia. Am J Psychiatry. 2003;160:13–23.

24. Buckner RL, Andrews-Hanna JR, Schacter DL. The Brain’s Default Network. Ann N Y Acad Sci. 2008;1124:1–38.

25. Zhou Y, Liang M, Tian L, Wang K, Hao Y, Liu H, et al. Functional disintegration in paranoid schizophrenia using resting-state fMRI. Schizophr Res. 2007;97:194–205.

26. Garrity AG, Pearlson GD, McKiernan K, Lloyd D, Kiehl KA, Calhoun VD. Aberrant ‘default mode’ functional connectivity in schizophrenia. Am J Psychiatry. 2007;164:450–457.

27. Harrison BJ, Yücel M, Pujol J, Pantelis C. Task-induced deactivation of midline cortical regions in schizophrenia assessed with fMRI. Schizophr Res. 2007;91:82–86.

28. Bressler SL, Menon V. Large-scale brain networks in cognition: emerging methods and principles. Trends Cogn Sci. 2010;14:277–290.

29. Hilker R, Helenius D, Fagerlund B, Skytthe A, Christensen K, Werge TM, et al. Heritability of Schizophrenia and Schizophrenia Spectrum Based on the Nationwide Danish Twin Register. Biol Psychiatry. 2018;83:492–498.

30. International Schizophrenia Consortium, Purcell SM, Wray NR, Stone JL, Visscher PM, O’Donovan MC, et al. Common polygenic variation contributes to risk of schizophrenia and bipolar disorder. Nature. 2009;460:748–752.

31. Schizophrenia Working Group of the Psychiatric Genomics Consortium. Biological insights from 108 schizophrenia-associated genetic loci. Nature. 2014;511:421–427.

32. Van der Auwera S, Wittfeld K, Shumskaya E, Bralten J, Zwiers MP, Onnink AMH, et al. Predicting brain structure in population-based samples with biologically informed genetic scores for schizophrenia. Am J Med Genet Part B Neuropsychiatr Genet Off Publ Int Soc Psychiatr Genet. 2017;174:324–332.

33. Van der Auwera S, Wittfeld K, Homuth G, Teumer A, Hegenscheid K, Grabe HJ. No association between polygenic risk for schizophrenia and brain volume in the general population. Biol Psychiatry. 2015;78:e41–42.

34. Alloza C, Cox SR, Blesa Cábez M, Redmond P, Whalley HC, Ritchie SJ, et al. Polygenic risk score for schizophrenia and structural brain connectivity in older age: A longitudinal connectome and tractography study. NeuroImage. 2018. 1 September 2018. https://doi.org/10.1016/j.neuroimage.2018.08.075.

35. McIntosh AM, Gow A, Luciano M, Davies G, Liewald DC, Harris SE, et al. Polygenic risk for schizophrenia is associated with cognitive change between childhood and old age. Biol Psychiatry. 2013;73:938–943.

36. Neilson E, Shen X, Cox SR, Clarke T-K, Wigmore EM, Gibson J, et al. Impact of Polygenic Risk for Schizophrenia on Cortical Structure in UK Biobank. Biol Psychiatry. 2019;0.

37. Ritchie SJ, Tucker-Drob EM, Cox SR, Dickie DA, Hernández M del CV, Corley J, et al. Risk and protective factors for structural brain ageing in the eighth decade of life. Brain Struct Funct. 2017:1–14.

38. Cox SR, Bastin ME, Ferguson KJ, Allerhand M, Royle NA, Maniega SM, et al. Compensation or inhibitory failure? Testing hypotheses of age-related right frontal lobe involvement in verbal memory ability using structural and diffusion MRI. Cortex J Devoted Study Nerv Syst Behav. 2015;63:4–15.

39. Hoffman P, Cox SR, Dykiert D, Muñoz Maniega S, Valdés Hernández MC, Bastin ME, et al. Brain grey and white matter predictors of verbal ability traits in older age: The Lothian Birth Cohort 1936. NeuroImage. 2017;156:394–402.

40. Kirkpatrick B, Messias E, Harvey PD, Fernandez-Egea E, Bowie CR. Is Schizophrenia a Syndrome of Accelerated Aging? Schizophr Bull. 2008;34:1024–1032.

41. Kochunov P, Glahn DC, Rowland LM, Olvera RL, Winkler A, Yang Y-H, et al. Testing the hypothesis of accelerated cerebral white matter aging in schizophrenia and major depression. Biol Psychiatry. 2013;73:482–491.

42. Hagenaars SP, Harris SE, Davies G, Hill WD, Liewald DCM, Ritchie SJ, et al. Shared genetic aetiology between cognitive functions and physical and mental health in UK Biobank (*N*=112 151) and 24 GWAS consortia. Mol Psychiatry. 2016;21:1624–1632.

43. Euesden J, Lewis CM, O’Reilly PF. PRSice: Polygenic Risk Score software. Bioinforma Oxf Engl. 2015;31:1466–1468.

44. Miller KL, Alfaro-Almagro F, Bangerter NK, Thomas DL, Yacoub E, Xu J, et al. Multimodal population brain imaging in the UK Biobank prospective epidemiological study. Nat Neurosci. 2016;19:1523–1536.

45. Desikan RS, Ségonne F, Fischl B, Quinn BT, Dickerson BC, Blacker D, et al. An automated labeling system for subdividing the human cerebral cortex on MRI scans into gyral based regions of interest. NeuroImage. 2006;31:968–980.

46. Roberts JA, Perry A, Roberts G, Mitchell PB, Breakspear M. Consistency-based thresholding of the human connectome. NeuroImage. 2017;145:118–129.

47. Menon V, Uddin LQ. Saliency, switching, attention and control: a network model of insula function. Brain Struct Funct. 2010;214:655–667.

48. Catani M, Dell’acqua F, Thiebaut de Schotten M. A revised limbic system model for memory, emotion and behaviour. Neurosci Biobehav Rev. 2013;37:1724–1737.

49. Power JD, Cohen AL, Nelson SM, Wig GS, Barnes KA, Church JA, et al. Functional network organization of the human brain. Neuron. 2011;72:665–678.

50. Sridharan D, Levitin DJ, Menon V. A critical role for the right fronto-insular cortex in switching between central-executive and default-mode networks. Proc Natl Acad Sci U S A. 2008;105:12569–12574.

51. Tucker-Drob EM. How Many Pathways Underlie Socioeconomic Differences in the Development of Cognition and Achievement? Learn Individ Differ. 2013;25:12–20.

52. Jöreskog KG, Goldberger AS. Estimation of a Model with Multiple Indicators and Multiple Causes of a Single Latent Variable. J Am Stat Assoc. 1975;70:631–639.

53. Rosseel Y. lavaan: An R Package for Structural Equation Modeling. J Stat Softw. 2012;48:36.

54. Ho D, Imai K, King G, Stuart EA. MatchIt: Nonparametric Preprocessing for Parametric Causal Inference. J Stat Softw. 2011;42:1–28.

55. Allen M, Poggiali D, Whitaker K, Marshall TR, Kievit R. Raincloud plots: a multi-platform tool for robust data visualization. PeerJ Inc.; 2018.

56. Nuevo R, Os JV, Arango C, Chatterji S, Ayuso-Mateos JL. Evidence for the early clinical relevance of hallucinatory-delusional states in the general population. Acta Psychiatr Scand. 2013. https://onlinelibrary.wiley.com/doi/abs/10.1111/acps.12010. Accessed 24 July 2019.

57. Benjamini Y, Hochberg Y. Controlling the False Discovery Rate: A Practical and Powerful Approach to Multiple Testing. J R Stat Soc Ser B Methodol. 1995;57:289–300.

58. Muñoz Maniega S, Lymer GKS, Bastin ME, Marjoram D, Job DE, Moorhead TWJ, et al. A diffusion tensor MRI study of white matter integrity in subjects at high genetic risk of schizophrenia. Schizophr Res. 2008;106:132–139.

59. Reus LM, Shen X, Gibson J, Wigmore E, Ligthart L, Adams MJ, et al. Association of polygenic risk for major psychiatric illness with subcortical volumes and white matter integrity in UK Biobank. Sci Rep. 2017;7.

60. Terwisscha van Scheltinga A, Bakker SC, van Haren NEM, Derks EM, Buizer-Voskamp JE, Boos HBM, et al. Genetic schizophrenia risk variants jointly modulate total brain and white matter volume. Biol Psychiatry. 2013;73:525–531.

61. Hulshoff Pol HE, Brans RGH, van Haren NEM, Schnack HG, Langen M, Baaré WFC, et al. Gray and white matter volume abnormalities in monozygotic and same-gender dizygotic twins discordant for schizophrenia. Biol Psychiatry. 2004;55:126–130.

62. Davis KAS, Cullen B, Adams M, Brailean A, Breen G, Coleman JRI, et al. Indicators of mental disorders in UK Biobank-A comparison of approaches. Int J Methods Psychiatr Res. 2019:e1796.

63. McGrath JJ, Saha S, Al-Hamzawi A, Alonso J, Bromet EJ, Bruffaerts R, et al. Psychotic Experiences in the General Population: A Cross-National Analysis Based on 31,261 Respondents From 18 Countries. JAMA Psychiatry. 2015;72:697–705.

64. Nuevo R, Chatterji S, Verdes E, Naidoo N, Arango C, Ayuso-Mateos JL. The continuum of psychotic symptoms in the general population: a cross-national study. Schizophr Bull. 2012;38:475–485.

65. Jones HJ, Stergiakouli E, Tansey KE, Hubbard L, Heron J, Cannon M, et al. Phenotypic Manifestation of Genetic Risk for Schizophrenia During Adolescence in the General Population. JAMA Psychiatry. 2016;73:221–228.

66. Zammit S, Hamshere M, Dwyer S, Georgiva L, Timpson N, Moskvina V, et al. A population-based study of genetic variation and psychotic experiences in adolescents. Schizophr Bull. 2014;40:1254–1262.

67. Critchley HD, Wiens S, Rotshtein P, Ohman A, Dolan RJ. Neural systems supporting interoceptive awareness. Nat Neurosci. 2004;7:189–195.

68. Damasio A. Mental self: The person within. Nature. 2003;423:227.

69. Fonville L, Cohen Kadosh K, Drakesmith M, Dutt A, Zammit S, Mollon J, et al. Psychotic Experiences, Working Memory, and the Developing Brain: A Multimodal Neuroimaging Study. Cereb Cortex N Y N 1991. 2015;25:4828–4838.

70. Crespo-Facorro B, Kim J, Andreasen NC, O’Leary DS, Bockholt HJ, Magnotta V. Insular cortex abnormalities in schizophrenia: a structural magnetic resonance imaging study of first-episode patients. Schizophr Res. 2000;46:35–43.

71. Díaz-Caneja CM, Schnack H, Martínez K, Santonja J, Alemán-Gomez Y, Pina-Camacho L, et al. Neuroanatomical deficits shared by youth with autism spectrum disorders and psychotic disorders. Hum Brain Mapp. 2019;40:1643–1653.

72. Crespo-Facorro B, Roiz-Santiáñez R, Quintero C, Pérez-Iglesias R, Tordesillas-Gutiérrez D, Mata I, et al. Insular cortex morphometry in first-episode schizophrenia-spectrum patients: Diagnostic specificity and clinical correlations. J Psychiatr Res. 2010;44:314–320.

73. Sommer IEC, Diederen KMJ, Blom J-D, Willems A, Kushan L, Slotema K, et al. Auditory verbal hallucinations predominantly activate the right inferior frontal area. Brain. 2008;131:3169–3177.

74. Francis AN, Seidman LJ, Jabbar GA, Mesholam-Gately R, Thermenos HW, Juelich R, et al. Alterations in brain structures underlying language function in young adults at high familial risk for schizophrenia. Schizophr Res. 2012;141:65–71.

75. Padmanabhan JL, Tandon N, Haller CS, Mathew IT, Eack SM, Clementz BA, et al. Correlations Between Brain Structure and Symptom Dimensions of Psychosis in Schizophrenia, Schizoaffective, and Psychotic Bipolar I Disorders. Schizophr Bull. 2015;41:154–162.

76. Paulesu E, Frith CD, Frackowiak RS. The neural correlates of the verbal component of working memory. Nature. 1993;362:342–345.

77. Gaser C, Nenadic I, Volz H-P, Büchel C, Sauer H. Neuroanatomy of ‘hearing voices’: a frontotemporal brain structural abnormality associated with auditory hallucinations in schizophrenia. Cereb Cortex N Y N 1991. 2004;14:91–96.

78. Hoffman RE, Rapaport J, Mazure CM, Quinlan DM. Selective speech perception alterations in schizophrenic patients reporting hallucinated ‘voices’. Am J Psychiatry. 1999;156:393–399.

79. Zmigrod L, Garrison JR, Carr J, Simons JS. The neural mechanisms of hallucinations: A quantitative meta-analysis of neuroimaging studies. Neurosci Biobehav Rev. 2016;69:113–123.

80. Corlett PR, Horga G, Fletcher PC, Alderson-Day B, Schmack K, Powers AR. Hallucinations and Strong Priors. Trends Cogn Sci. 2019;23:114–127.

81. Bohland JW, Bokil H, Allen CB, Mitra PP. The brain atlas concordance problem: quantitative comparison of anatomical parcellations. PloS One. 2009;4:e7200.

82. Cox SR, Ferguson KJ, Royle NA, Shenkin SD, MacPherson SE, MacLullich AMJ, et al. A systematic review of brain frontal lobe parcellation techniques in magnetic resonance imaging. Brain Struct Funct. 2014;219:1–22.

83. de Reus MA, van den Heuvel MP. The parcellation-based connectome: limitations and extensions. NeuroImage. 2013;80:397–404.

84. Goulas A, Uylings HBM, Hilgetag CC. Principles of ipsilateral and contralateral cortico-cortical connectivity in the mouse. Brain Struct Funct. 2017;222:1281–1295.

85. Buchanan CR, Bastin ME, Ritchie SJ, Liewald DC, Madole J, Tucker-Drob EM, et al. The effect of network thresholding and weighting on structural brain networks in the UK Biobank. BioRxiv. 2019. 24 May 2019. https://doi.org/10.1101/649418.

86. Fry A, Littlejohns TJ, Sudlow C, Doherty N, Adamska L, Sprosen T, et al. Comparison of Sociodemographic and Health-Related Characteristics of UK Biobank Participants With Those of the General Population. Am J Epidemiol. 2017;186:1026–1034.

