## Supplementary material for "Psychotic-like experiences, polygenic risk scores for schizophrenia and structural properties of the salience, default mode and central-executive networks in healthy participants from UK Biobank": SM

**Supplementary Material Table 1: Psychiatric diagnoses**

A total of 163 participants with a psychiatric-related admission were excluded from analyses. Diagnoses of mental and behavioural disorders were coded according to the International Classification of Disease version 10 (ICD10). Due to comorbidity, the total number of diagnoses is 244 for each individual (including those that were initially excluded due to a diagnosis of schizophrenia or bipolar disorder with psychotic symptoms).

| Diagnosis | IC10 code | N |
|---|---|---|
| Mild cognitive disorder | F067 | 1 |
| Postconcussional disorder | F072 | 1 |
| Acute intoxication | F100 | 8 |
| Harmful use - Alcohol | F101 | 11 |
| Dependence syndrome - Alcohol | F102 | 6 |
| Withdrawal state - Alcohol | F103 | 5 |
| Amnesic syndrome - Alcohol | F106 | 2 |
| Dependence syndrome - Opioids | F112 | 2 |
| Harmful use - Tobacco | F171 | 39 |
| Dependence syndrome - Tobacco | F172 | 1 |
| Psychotic disorder - Drug use | F195 | 1 |
| Schizophrenia, unspecified | F209 | 3 |
| Delusional disorder | F220 | 1 |
| Schizoaffective disorder, unspecified | F259 | 2 |
| Unspecified nonorganic psychosis | F29 | 3 |
| Bipolar affective disorder, unspecified | F319 | 3 |
| Mild depressive episode | F320 | 2 |
| Severe depressive episode without psychotic symptoms | F322 | 4 |
| Severe depressive episode with psychotic symptoms | F323 | 2 |
| Depressive episode, unspecified | F329 | 84 |
| Recurrent depressive disorder, current episode mild | F330 | 1 |
| Recurrent depressive disorder, current episode moderate | F331 | 2 |
| Recurrent depressive disorder, current episode severe without psychotic symptoms | F332 | 1 |
| Recurrent depressive disorder, unspecified | F339 | 4 |
| Dysthymia | F341 | 1 |
| Unspecified mood [affective] disorder | F39 | 1 |
| Agoraphobia | F400 | 1 |

| | | |
|---|---|---|
| Specific (isolated) phobias | F402 | 4 |
| Other phobic anxiety disorders | F408 | 1 |
| Panic disorder [episodic paroxysmal anxiety] | F410 | 2 |
| Mixed anxiety and depressive disorder | F412 | 5 |
| Anxiety disorder, unspecified | F419 | 21 |
| Obsessive-compulsive disorder, unspecified | F429 | 1 |
| Acute stress reaction | F430 | 1 |
| Adjustment disorders | F432 | 2 |
| Reaction to severe stress, unspecified | F439 | 1 |
| Dissociative motor disorders | F444 | 1 |
| Somatoform autonomic dysfunction | F453 | 1 |
| Other somatoform disorders | F458 | 4 |
| Neurasthenia | F480 | 1 |
| Neurotic disorder, unspecified | F489 | 1 |
| Failure of genital response | F522 | 1 |
| Severe mental and behavioural disorders associated with the puerperium, not elsewhere classified | F531 | 2 |
| Emotionally unstable personality disorder | F603 | 1 |
| Anankastic personality disorder | F605 | 1 |
| Asperger's syndrome | F845 | 1 |

**Supplementary Material Figure 1: Diagram of MRI data used in this study.**

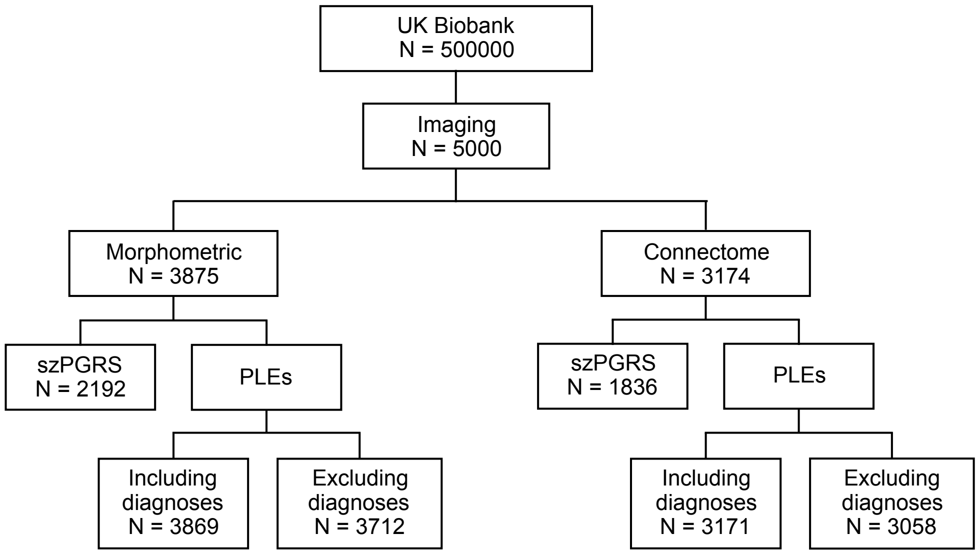

*Note:* $PRS_{SZ}$: polygenic risk for schizophrenia; PLEs: Psychotic-like experiences. Valid sample sizes for PLEs analyses after matching subjects 1:10 are presented in Table 2 in the main manuscript.

**Supplementary Material Methods 1**

**Scan acquisition**

MRI data for all participants were acquired on a single Siemens Skyra 3 T scanner. Briefly, the acquired 3D MPRAGE T1-weighted volumes were pre-processed and analyzed using FSL tools (http://www.fmrib.ox.ac.uk/fsl) by the UK Biobank brain imaging team. This included a raw, de-faced T1-weighted volume, a reduced field-of-view (FoV) T1-weighted volume, and further processing, which included skull stripping, bias field correction and gross tissue segmentation using FNIRT [1] and FAST [2], yielding cerebrospinal fluid (CSF), grey and white matter volumes. Where large, common artefacts, such as head movement, were identified during scanning, image acquisition was re-started. However, visual quality control was not systematically undertaken by the UK Biobank team, as this would be impractical due to the very large sample size [3]. No significant changes were made to scanner hardware or software during the period of MRI data acquisition; full details on protocol phases and relevant upgrades are available at the following URL: http://biobank.ctsu.ox.ac.uk/crystal/docs/brain_mri.pdf.

**Local Image Processing**

We undertook further image processing locally, using bulk image data downloaded from UK Biobank. Each 3D $T_1$-weighted FSPGR volume was parcellated into 85 regions-of-interest (ROI) using FreeSurfer version 5.3 (which comprised 34 cortical ROIs and eight sub-cortical ROIs per hemisphere, plus the brainstem). Segmentations were visually checked and removed case-wise in instances of aberrant surfaces, tissue identification or segmentation failure, due to the characteristics of the sample and analyses no outliers were excluded. The outputs were then used to construct grey and white matter masks for use in network construction and to constrain the tractography output as described below. According to previously published methods [4], we used tools provided by the FDT package in FSL (http://fsl.fmrib.ox.ac.uk/fsl) to pre-process the diffusion MRI data, reducing systematic imaging distortions and bulk subject motion artefacts by affine registration of all subsequent echo-planar (EP) volumes to the first $T_2$-weighted EP volume [5]. Skull stripping and brain extraction were performed on the registered $T_2$-weighted EP volumes and applied to the mean diffusivity (MD)/FA volumes calculated by DTIFIT in each subject [6, 7]. The neuroanatomical ROIs determined by FreeSurfer were then aligned from 3D $T_1$-weighted volume to diffusion space using a cross-modal nonlinear registration method. As a first step, linear registration was used to initialize the alignment of each brain-extracted FA volume to the corresponding FreeSurfer extracted 3D $T_1$-weighted brain volume using a mutual information cost function and an affine transform with 12 degrees of freedom [5]. Following this initialization, a nonlinear deformation field based method (FNIRT) was used to refine local alignment [1]. FreeSurfer segmentations and anatomical labels were then aligned to diffusion space using nearest neighbour interpolation. Whole-brain probabilistic tractography was performed using FSL's BedpostX/ProbTrackX algorithm [8]. Probability density functions, which describe the uncertainty in the principal directions of water diffusion, were computed using a two-fibre model per voxel [8]. Streamlines were then constructed by sampling from these distributions during a tracking process that involved all white matter voxels using 100 Markov Chain Monte Carlo iterations with a fixed step size of 0.5 mm between successive points. Tracking was initiated from all white matter voxels [4] in two collinear directions until terminated by the following stopping criteria designed to minimize the amount of anatomically implausible streamlines: (i) exceeding a curvature threshold of 70 degrees; (ii) entering a voxel with FA below 0.1 ; (iii) entering an extra-cerebral voxel; (iv) exceeding 200 mm in length; and (v) exceeding a distance ratio metric of 10. The distance ratio metric [9], excludes implausibly tortuous streamlines. For instance, a streamline with a total path length 10 times longer than the distance between end points was considered to be invalid. The values of the curvature, anisotropy and distance ratio metric constraints were set empirically and informed by visual assessment of the resulting streamlines.

**Supplementary Material Table 2: Desikan-Killiany Abbreviations**

| Desikan-Killiany | Abbreviation | Desikan-Killiany (cont.) | Abbreviation (cont.) |
|---|---|---|---|
| Left-Thalamus-Proper | L-Thal | ctx-lh-superiorfrontal | L-Sup Front |
| Left-Caudate | L-Caudate | ctx-lh-superiorparietal | L-Sup Par |
| Left-Putamen | L-Putamen | ctx-lh-superiortemporal | L-Sup Temp |
| Left-Pallidum | L-Pallidum | ctx-lh-supramarginal | L-Supramar |
| Brain-Stem | BS | ctx-lh-frontalpole | L-FrontPole |
| Left-Hippocampus | L-Hipp | ctx-lh-temporalpole | L-TempPole |
| Left-Amygdala | L-Amyg | ctx-lh-transversetemporal | L-TransT |
| Left-Accumbens-area | L-Acc | ctx-lh-insula | L-Insula |
| Left-VentralDC | L-VentralDC | ctx-rh-bankssts | R-Bankssts |
| Right-Thalamus-Proper | R-Thal | ctx-rh-caudalanteriorcingulate | R-CAC |
| Right-Caudate | R-Caudate | ctx-rh-caudalmiddlefrontal | R-Cau Mid F |
| Right-Putamen | R-Putamen | ctx-rh-cuneus | R-Cuneus |
| Right-Pallidum | R-Pallidum | ctx-rh-entorhinal | R-Entorhinal |
| Right-Hippocampus | R-Hippocampus | ctx-rh-fusiform | R-Fusiform |
| Right-Amygdala | R-Amyg | ctx-rh-inferiorparietal | R-Inf Par |
| Right-Accumbens-area | R-Acc | ctx-rh-inferiortemporal | R-Inf Temp |
| Right-VentralDC | R-VentralDC | ctx-rh-isthmuscingulate | R-Isthmus |
| ctx-lh-bankssts | L-Bankssts | ctx-rh-lateraloccipital | R-Lat Occ |
| ctx-lh-caudalanteriorcingulate | L-CAC | ctx-rh-lateralorbitofrontal | R-Lat Orb F |
| ctx-lh-caudalmiddlefrontal | L-Cau Mid F | ctx-rh-lingual | R-lingual |
| ctx-lh-cuneus | L-Cuneus | ctx-rh-medialorbitofrontal | R-Med OrbF |
| ctx-lh-entorhinal | L-Entorhinal | ctx-rh-middletemporal | R-Midd Temp |
| ctx-lh-fusiform | L-Fusiform | ctx-rh-parahippocampal | R-ParaHipp |
| ctx-lh-inferiorparietal | L-Inf Par | ctx-rh-paracentral | R-ParaCent |
| ctx-lh-inferiortemporal | L-Inf Temp | ctx-rh-parsopercularis | R-Operc |
| ctx-lh-isthmuscingulate | L-Isthmus | ctx-rh-parsorbitalis | R-Orbitalis |
| ctx-lh-lateraloccipital | L-Lat Occ | ctx-rh-parstriangularis | R-Triang |
| ctx-lh-lateralorbitofrontal | L-Lat Orb F | ctx-rh-pericalcarine | R-Perical |
| ctx-lh-lingual | L-lingual | ctx-rh-postcentral | R-PostCentr |
| ctx-lh-medialorbitofrontal | L-Med OrbF | ctx-rh-posteriorcingulate | R-Post Cing |
| ctx-lh-middletemporal | L-Midd Temp | ctx-rh-precentral | R-PreCent |
| ctx-lh-parahippocampal | L-ParaHipp | ctx-rh-precuneus | R-Prec |
| ctx-lh-paracentral | L-ParaCent | ctx-rh-rostralanteriorcingulate | R-RAC |
| ctx-lh-parsopercularis | L-Operc | ctx-rh-rostralmiddlefrontal | R-Rost Midd F |
| ctx-lh-parsorbitalis | L-Orbitalis | ctx-rh-superiorfrontal | R-Sup Front |
| ctx-lh-parstriangularis | L-Triang | ctx-rh-superiorparietal | R-Sup Par |
| ctx-lh-pericalcarine | L-Perical | ctx-rh-superiortemporal | R-Sup Temp |
| ctx-lh-postcentral | L-PostCentr | ctx-rh-supramarginal | R-Supramar |
| ctx-lh-posteriorcingulate | L-Post Cing | ctx-rh-frontalpole | R-FrontPole |
| ctx-lh-precentral | L-PreCent | ctx-rh-temporalpole | R-TempPole |

| | | | |
|---|---|---|---|
| ctx-lh-precuneus | L-Prec | ctx-rh-transversetemporal | R-TransT |
| ctx-lh-rostralanteriorcingulate | L-RAC | ctx-rh-insula | R-Insula |
| ctx-lh-rostralmiddlefrontal | L-Rost Midd F | | |

**Supplementary Material Results 1: $PRS_{SZ}$ at p ≤ 0.5 and p ≤ 1.0 thresholds**

**$PRS_{SZ}$ analyses**

**Salience Network**

*Linear regressions for individual network components*

There were no significant associations between cortical and subcortical volume and $PRS_{SZ}$ at any thresholds ($p_{FDR} > 0.05$). However, for cortical thickness, significant negative associations were found between the right insula and $PRS_{SZ}$ at a threshold of p ≤ 0.5 (β = -0.042, $p = 0.026$) and p ≤ 1.0 (β = -0.038, $p = 0.042$) although none of these associations survived FDR-correction ($p_{FDR} > 0.05$). Interactions between age and $PRS_{SZ}$ were not significant for cortical thickness or grey matter volume ($p_{FDR} > 0.05$). There were no significant associations between white matter FA and $PRS_{SZ}$ at any threshold or with the interaction between age and $PRS_{SZ}$ ($p_{FDR} > 0.05$).

*MIMIC common + independent pathways analysis*

The associations between the latent factor for grey matter volume and $PRS_{SZ}$ were not significant at any threshold (p ≤ 0.5: r = -0.027, SE = 0.013; p ≤ 1.0: r = -0.020, SE = 0.013; all $p_{FDR} > 0.05$). Allowing for a direct effect of $PRS_{SZ}$ on left thalamus volume significantly improved model fit at all levels of polygenic significance thresholding ($\chi^2[1] > 4.917$, p < 0.028; independent pathway estimates: p ≤ 0.5: β = 0.022, $p = 0.006$; p ≤ 1.0: β = 0.022, $p = 0.006$).

There were no significant associations between the latent factor for FA and $PRS_{SZ}$ at any threshold (all thresholds: r = 0.004, SE = 0.017; all $p_{FDR} > 0.05$). An independent pathway emerged between $PRS_{SZ}$ and the white matter connection between the left thalamus and left ventral diencephalon, though this effect was only significant at p ≤ 0.5 and p ≤ 1.0 of polygenic thresholding ($\chi^2[1] > 5.568$, $p < 0.019$; independent pathway estimates: both p ≤ 0.5 and p ≤ 1.0: β = 0.051, $p$ =0.017). Interactions between $PRS_{SZ}$*age were not significantly associated with any latent construct in the salience network ($p_{FDR}$ >

0.05).

**Default Mode Network**

*Linear regressions for individual network components*

For cortical thickness, significant negative associations were found between $PRS_{SZ}$ at a threshold of $p \leq 1.0$ and the left latero-orbito frontal ($\beta = -0.039$, $p = 0.044$), and right medialorbito frontal ($\beta = -0.046$, $p = 0.037$). For grey matter volume, significant negative associations were found between $PRS_{SZ}$ at the thresholds of $p \leq 0.5$, $p \leq 1.0$ and the left latero-orbito frontal ($\beta_{p \leq 0.5} = -0.028$, $p_{p \leq 0.5} = 0.048$; $\beta_{p \leq 1.0} = -0.028$, $p_{p \leq 1.0} = 0.045$), and right medialorbito frontal ($\beta_{p \leq 0.5} = -0.033$, $p_{p \leq 0.5} = 0.043$; $\beta_{p \leq 1.0} = -0.033$, $p_{p \leq 1.0} = 0.045$). Although none of these associations survived FDR-correction ($p_{FDR} > 0.05$). There were no significant associations FA and $PRS_{SZ}$ or the interaction between $PRS_{SZ}$*age at any thresholds ($p_{FDR} > 0.05$).

*MIMIC common + independent pathways analysis*

The associations between the latent factor for grey matter volume and $PRS_{SZ}$ were significant $p \leq 0.5$ ($p \leq 0.5$: r = -0.047, SE = 0.014, $p = 0.043$ ; $p \leq 1.0$: r = -0.043, SE = 0.014, $p = 0.065$). Albeit none of these associations survived multiple comparison correction, the directions and magnitudes of all associations were consistent ($p_{FDR} > 0.05$). Allowing a direct effect of $PRS_{SZ}$ on right pars orbitalis volume significantly improved model fit, though this was only found at $p \leq 0.5$ of polygenic thresholding ($\chi^2[1] = 4.112$, $p = 0.043$; independent pathway estimate: $\beta = 0.033$, $p = 0.043$).

Associations between the latent factor for grey matter thickness and $PRS_{SZ}$ were not significant at any threshold ($p \leq 0.5$: r = -0.030, SE = 0.015; $p \leq 1.0$: r = -0.034, SE = 0.015; all $p_{FDR} > 0.05$). Age showed a significant positive effect on the latent factor (r = 0.210, SE = 0.017, $p < 0.001$). An independent pathway emerged from $PRS_{SZ}$*age to right supramarginal thickness at all levels of polygenic significance thresholding ($\chi^2[1] > 5.705$, $p < 0.018$; all independent pathway estimates: $p \leq 0.5$ and $p \leq 1.0$: $\beta = -0.035$, $p = 0.008$). At $p \leq 0.5$ and $p \leq 1.0$, a second direct effect of $PRS_{SZ}$*age on left posterior cingulate thickness significantly improved model fit relative to the model including only the direct effect on right supramarginal thickness ($\chi^2[1] > 5.440$, $p < 0.021$; both independent pathway estimates:

$p \leq 0.5$ and ; $p \leq 1.0$: $\beta = -0.043$, $p = 0.018$). Interactions between $PRS_{SZ}$*age were not significantly associated with any latent construct in the DMN ($p_{FDR} > 0.05$).

#### Central Executive Network

*Linear regressions for individual network components*

A significant positive association was found between the volume of the right inferior parietal and $PRS_{SZ}$ at a threshold of $p \leq 0.5$ ($\beta = 0.047$, SE = 0.015, $p_{FDR} = 0.027$) and $p \leq 1.0$ ($\beta = 0.046$, SE = 0.015, $p_{FDR} = 0.017$). A significant positive association was found between $PRS_{SZ}$ at a threshold of $p \leq 1.0$ and FA connecting the right inferior parietal with the right supramarginal ($\beta = 0.048$, $p = 0.038$) but it did not survive FDR-correction ($p_{FDR} > 0.05$). No significant associations were found for the other brain metrics of the CEN and $PRS_{SZ}$ ($p_{FDR} > 0.05$). The interaction between $PRS_{SZ}$*age was not significant at any thresholds ($p_{FDR} > 0.05$).

*MIMIC common + independent pathways analysis*

The associations between the latent factors for grey matter and $PRS_{SZ}$ were not significant at any threshold (volume: $p \leq 0.5$: r = 0.001, SE = 0.009; $p \leq 1.0$: r = 0.012, SE = 0.009, all $p_{FDR} > 0.05$; age $_{volume}$ r = -0.315, SE = 0.011, $p < 0.001$; cortical thickness: $p \leq 0.5$: r = 0.041, SE = 0.010; $p \leq 1.0$: r = 0.041, SE = 0.010, all $p_{FDR} > 0.05$; age $_{thickness}$ r = 0.083, SE = 0.012, $p = 0.006$). For the volumetric model, allowing for a direct effect of $PRS_{SZ}$ on right inferior parietal volume resulted in significant improvement in model fit at all levels of polygenic significance thresholding ($\chi^2[1] > 7.741$, $p < 0.006$; independent pathway estimates: $p \leq 0.5$: $\beta = 0.046$, $p = 0.001$; $p \leq 1.0$: $\beta = 0.047$, $p = 0.001$). For the model assessing cortical thickness, an independent pathway emerged from $PRS_{SZ}$*age to right superior frontal thickness at all levels of polygenic significance thresholding ($\chi^2[1] > 7.512$, $p < 0.007$; independent pathway estimates: $p \leq 0.5$ and $p \leq 1.0$: $\beta = 0.028$, $p = 0.002$). Adding a second direct effect from $PRS_{SZ}$*age to right inferior parietal thickness significantly improved model fit at all levels of polygenic significance relative to the model including only the direct effect on right superior frontal thickness ($\chi^2[1] > 4.314$, $p < 0.039$; independent pathway estimates: $p \leq 0.5$ and $p \leq 1.0$: $\beta = -0.024$, $p = 0.034$). Interactions between $PRS_{SZ}$*age were not significantly associated with any latent construct in the CEN ($p_{FDR} > 0.05$).

**Supplementary Material Methods 2: MIMIC common + independent pathways analysis**

In this analysis, a common factor –for each NOI – was regressed on both $PRS_{SZ}$ and the interaction between $PRS_{SZ}$ and age simultaneously ("common pathway"), adjusting for age, sex and whole brain measures at the manifest level, model fits were assessed, and residual correlations were modelled as above. Direct effects from $PRS_{SZ}$ to the indicators of the common factors (i.e., nodes or edges) are then added one by one ("independent pathways"), beginning with the effect that produces the largest increment in model fit and stopping once addition of direct effects does not result in statistically significant improvements in model fit at $p < 0.050$. The addition of direct effects here would indicate that the effect of $PRS_{SZ}$ on brain network integrity may not act solely on a global network level but may be influenced by one or more constituent parts of the network (i.e nodes or white matter tracts). Significant differences in model fit were assessed using a chi-square difference test. With two exceptions (FA in the DMN and CEN, whose measurement models did not fit well: RMSEA > 0.050, CFI < 0.950, SRMR > 0.050), this analysis was run for all three factors of network integrity in each of the three networks studied. In those cases in which measurement models of global network integrity did not fit well (FA models within the DMN and CEN), this indicated that the association between these network components and $PRS_{SZ}$ was best interpreted as independent associations (i.e. as indicated by the initial linear regressions).

**Supplementary Material Results 2: PLEs in participants without any psychiatric diagnosis**

In order to determine whether the signal could be driven by any psychiatric disorder beyond a diagnosis of psychosis, we excluded all participants with a psychiatric status from the PLEs analysis (their diagnoses can be found in Supplementary Material Table 1). Within the salience network, significant negative associations were found between the thickness of the right insula and auditory hallucinations ($N_{cases} = 46$, $N_{controls} = 460$; $\beta = -0.110$, SE = 0.039, $p_{FDR} = 0.019$). Moreover, FA of pathway connecting the left insula with the left amygdala showed a tendency towards significance with persecutory delusions ($N_{cases} = 9$, $N_{controls} = 90$; $\beta = -0.282$, SE = 0.098, $p = 0.005$, $p_{FDR} = 0.070$). Within the DMN, we found a positive association between persecutory delusions and cortical thickness of the right supramarginal ($N_{cases}$ = 10, $N_{controls}$ = 100; β = 0.171, SE = 0.050, $p_{FDR}$ = 0.009), and between the volume of the right supramarginal and delusions of reference ($N_{cases}$ = 11, $N_{controls}$ = 110; β = - 0.227, SE = 0.067, $p_{FDR}$ = 0.014). By excluding participants with any psychiatric disorder, results from this analysis indicate that these significant associations between brain structure and PLEs represent a continuous trait in the general population which are not driven by any psychiatric diagnosis. This study also found a significantly negative association between total number of PLEs and the volume of the right medial orbito frontal exclusively in participants without any psychiatric diagnosis, suggesting that this node may present structural alterations in a non-symptom-specific manner. From this data, we can conclude that certain associations between neurostructural properties and PLEs are present in the general population which are not driven by any psychiatric diagnosis, while those associations are enriched for psychiatric cases.

**Supplementary Material Results 3: Level distress in relation to PLEs**

We tested whether distress was significantly associated with PLEs using logistic regression controlling for age and sex. Level of distress in relation to PLEs was categorised by "Not distressing at all, it was a positive experience", "Not distressing, a neutral experience", "A bit distressing", "Quite distressing", and "Very distressing". We found that persecutory delusions were significantly associated with level of distress (β = 1.68, SE = 0.388, $p < 0.001$). Thus, we tested whether the associations found between our nodes and persecutory delusions were significantly mediated by level of distress. However, we did not find any significant mediation effect (*p* values < 0.05).

**Supplementary Material Results 4: Level distress in relation to total number of PLEs**

We tested whether distress was significantly associated with the total sum of PLEs using general linear regression controlling for age and sex, and level of distress as the dependent variable. Level of distress in relation to PLEs was categorised by "Not distressing at all, it was a positive experience", "Not distressing, a neutral experience", "A bit distressing", "Quite distressing", and "Very distressing". We found that increased number of PLEs was significantly associated with higher level of distress (β = 0.093, SE = 0.039, $p$ = 0.020). Age also showed a significant effect (β = -0.215, SE = 0.081, $p$ = 0.009). This is in line with previous research [10], suggesting that co-occurrence of psychotic symptoms in the general population may be more clinically relevant.

**Supplementary Material Table 3: Participants with and without PLEs**

Based on participants answering either "Yes" or "No" to PLEs, the prevalence of at least one PLEs in this sample was 4.75% (N = 2819). Sample characteristics for participants reporting PLEs versus non-PLEs participants were broadly similar:

| Variable | Units | N | No PLEs | N | Any PLEs |
|---|---|---|---|---|---|
| **Age** | Years (SD) | 3595 | 62.24 (7.56) | 117 | 61.46 (7.83) |
| **Sex** | % Females | 3595 | 51.88% | 117 | 61.54% |
| **Grey matter cortical thickness** | mm Mean (SD) | 3330 | 2.40 (0.10) | 107 | 2.41 (0.10) |
| **Grey matter volume** | $mm^3$ Mean (SD) | 3589 | 620909.24 (55897.84) | 116 | 614557.03 (51204.20) |
| **Whole-brain FA** | Mean (SD) | 2956 | 0.46 (0.02) | 102 | 0.46 (0.01) |

**Supplementary Material Table 4: Networks fits and loadings at a $PRS_{SZ}$ threshold of $p \leq 0.1$**

| Network | RMSEA | CFI | SRMR | Standardised Loadings |
|---|---|---|---|---|
| **Salience** | | | | |
| Latent factor cortical thickness | 0.048 | 0.978 | 0.020 | 0.169 to 0.571 |
| Latent factor volume | 0.048 | 0.984 | 0.025 | -0.059 to 0.596 |
| Latent factor FA | 0.051 | 0.914 | 0.037 | -0.054 to 0.273 |
| **DMN** | | | | |
| Latent factor cortical thickness | 0.042 | 0.977 | 0.020 | -0.134 to 0.507 |
| Latent factor volume | 0.049 | 0.986 | 0.022 | 0.049 to 0.528 |
| Latent factor FA | 0.056 | 0.955 | 0.041 | 0.030 to 0.757 |
| **CEN** | | | | |
| Latent factor cortical thickness | 0.062 | 0.982 | 0.020 | -0.161 to 0.358 |
| Latent factor volume | 0.036 | 0.992 | 0.014 | -0.011 to 0.371 |
| Latent factor FA | 0.062 | 0.947 | 0.037 | 0.530 to 0.819 |

**Supplementary Material Figure 2: Mean cortical values for thickness and grey matter volume**

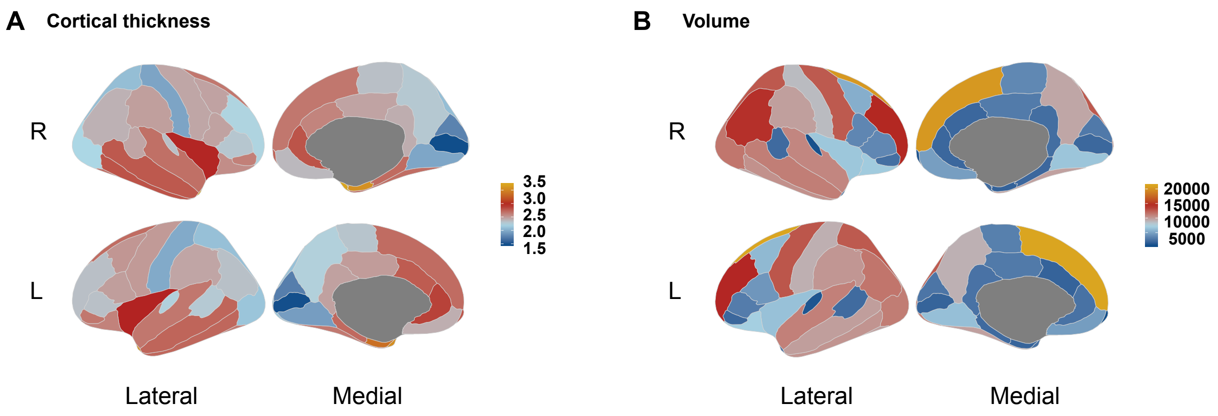

Lateral and medial views of **A** mean cortical values for grey matter thickness and **B** cortical volumes based on Desikan-Killiany atlas parcellation. A total sample of 3875 subjects was used to calculate the average morphometric properties. R = right hemisphere, L = left hemisphere, cortical thickness was measured in mm and cortical volumes in $mm^3$.

**Supplementary Material Table 5: Summary of associations between PLEs and brain structure**

| PLEs | Direction | Area | Metric | Network | $\beta_{standardised}$ | $P_{FDR}$ |
|---|---|---|---|---|---|---|
| **Auditory hallucinations** | - | Right Insula | Thickness | Salience | -0.114 | 0.004* |
| | - | Left Insula | Thickness | Salience | -0.083 | 0.045* |
| **Delusions of reference** | - | Right Supramarginal | Volume | DMN | -0.195 | 0.022* |
| **Persecutory delusions** | + | Right Supramarginal gyrus | Thickness | DMN | 0.177 | 0.003* |
| | + | Left Pars Orbitalis | Volume | DMN | 0.219 | 0.021* |

*Note:* Asterisks represent significant associations ($p_{FDR} < 0.050$).

**Supplementary Material Results 5: Mediation analyses at p ≤ 0.5 and p ≤ 1.0 $PRS_{SZ}$ thresholds**

Based on the significant associations between brain structure and PLEs in the salience and DMN, we examined the hypothesis that higher $PRS_{SZ}$ was associated with symptom severity via brain structure. First, we tested the association between $PRS_{SZ}$ and total number of symptoms in the whole UKBiobank sample, independently of diagnosis (N = 308693). However, we did not find any significant association ($p \leq 0.5$ : $\beta = 0.027$, SE = 0.018, $p = 0.143$; $p \leq 1.0$ : $\beta = 0.023$, SE = 0.018, $p = 0.201$). The mediation models using $PRS_{SZ}$ at 0.5 and 1.0 thresholds and mediators met the criteria for close model fit (RMSEA = 0, CFI = 1, SRMR < 0.05). Standardised parameter estimates for the model are presented in Figure 1. Our results show that the linear associations between $PRS_{SZ}$ and auditory hallucinations were significantly mediated by cortical thickness of the right insula at three different thresholds ($p \leq 0.5$: from $\sigma = 0.014$ to $\sigma' = -0.004$, CI [0.023, 0.133]; $p \leq 1.0$: from $\sigma = 0.016$ to $\sigma' = -0.002$, CI [0.017, 0.118], with the right insular cortex mediating 125% and 112% of the association between $PRS_{SZ}$ and auditory hallucinations, respectively). These results suggest that a negative association between $PRS_{SZ}$ and cortical thickness of the insula ($\varepsilon$) and a negative association between cortical thickness of the insula and auditory hallucinations ($\lambda$), results in a significant positive indirect association between $PRS_{SZ}$ and auditory hallucinations ($\sigma'$) with the inclusion of insular cortex as a mediator. No significant mediations were observed for any other ROIs.

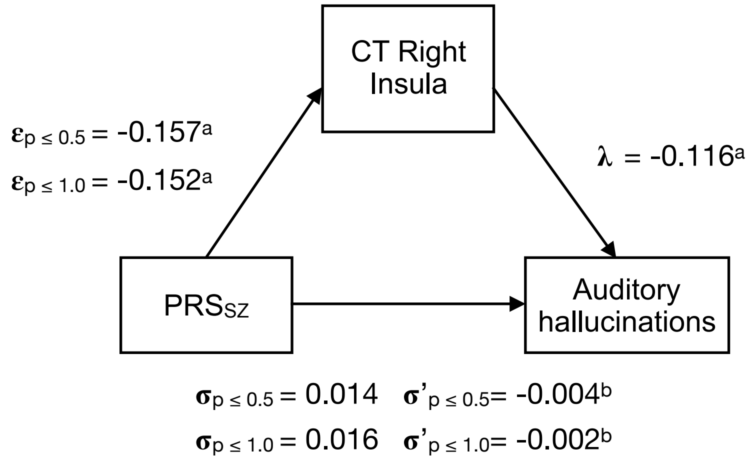

**Figure 1:** Path diagram of the mediation model, where the $\varepsilon$ coefficient represents the coefficients of regressions of $PRS_{SZ}$ at all thresholds on cortical thickness of the right insula, and $\lambda_{all}$ the coefficients of the regression of cortical thickness of the right insula on auditory hallucinations and $\sigma$ coefficients of the direct path of $PRS_{SZ}$ and auditory hallucinations at different $PRS_{SZ}$ thresholds. Coefficients of $\sigma'$ represents the indirect path of $PRS_{SZ}$ on auditory hallucinations through cortical thickness of the right

insula at different $PRS_{SZ}$ thresholds. *Note:* CT = cortical thickness, [a] $p$ values < 0.05, [b] Confidence intervals not including zero.

### References

1. Andersson JL, Jenkinson M, Smith S. Non-linear registration, aka Spatial normalisation FMRIB technical report TR07JA2. FMRIB Anal Group Univ Oxf. 2007;2.
2. Zhang Y, Brady M, Smith S. Segmentation of brain MR images through a hidden Markov random field model and the expectation-maximization algorithm. IEEE Trans Med Imaging. 2001;20:45–57.
3. Alfaro-Almagro F, Jenkinson M, Bangerter NK, Andersson JLR, Griffanti L, Douaud G, et al. Image processing and Quality Control for the first 10,000 brain imaging datasets from UK Biobank. NeuroImage. 2018;166:400–424.
4. Buchanan CR, Pernet CR, Gorgolewski KJ, Storkey AJ, Bastin ME. Test-retest reliability of structural brain networks from diffusion MRI. NeuroImage. 2014;86:231–243.
5. Jenkinson M, Smith S. A global optimisation method for robust affine registration of brain images. Med Image Anal. 2001;5:143–156.
6. Basser PJ, Pierpaoli C. Microstructural and physiological features of tissues elucidated by quantitative-diffusion-tensor MRI. J Magn Reson B. 1996;111:209–219.
7. Smith SM. Fast robust automated brain extraction. Hum Brain Mapp. 2002;17:143–155.
8. Behrens TEJ, Berg HJ, Jbabdi S, Rushworth MFS, Woolrich MW. Probabilistic diffusion tractography with multiple fibre orientations: What can we gain? NeuroImage. 2007;34:144–155.
9. Bullitt E, Gerig G, Pizer SM, Lin W, Aylward SR. Measuring tortuosity of the intracerebral vasculature from MRA images. IEEE Trans Med Imaging. 2003;22:1163–1171.
10. Nuevo R, Os JV, Arango C, Chatterji S, Ayuso-Mateos JL. Evidence for the early clinical relevance of hallucinatory-delusional states in the general population. Acta Psychiatr Scand. 2013. https://onlinelibrary.wiley.com/doi/abs/10.1111/acps.12010. Accessed 24 July 2019.
